# Ornithine capture by a translating ribosome controls bacterial polyamine synthesis

**DOI:** 10.1101/604074

**Authors:** Alba Herrero del Valle, Britta Seip, Iñaki Cervera-Marzal, Guénaël Sacheau, A. Carolin Seefeldt, C. Axel Innis

**Affiliations:** Institut Européen de Chimie et Biologie, Univ. Bordeaux, Institut National de la Santé et de la Recherche Médicale (U1212) and Centre National de la Recherche Scientifique (UMR 5320), Pessac 33607, France

## Abstract

Polyamines are essential metabolites that play an important role in cell growth, stress adaptation, and microbial virulence^1–3^. In order to survive and multiply within a human host, pathogenic bacteria adjust the expression and activity of polyamine biosynthetic enzymes in response to different environmental stresses and metabolic cues^2^. Here, we show that ornithine capture by the ribosome and the nascent peptide SpeFL controls polyamine synthesis in γ-proteobacteria by inducing the expression of the ornithine decarboxylase SpeF^4^, via a mechanism involving ribosome stalling and transcription antitermination. In addition, we present the cryo-EM structure of an *Escherichia coli* (*E. coli*) ribosome stalled during translation of *speFL* in the presence of ornithine. The structure shows how the ribosome and the SpeFL sensor domain form a highly selective binding pocket that accommodates a single ornithine molecule but excludes near-cognate ligands. Ornithine pre-associates with the ribosome and is then held in place by the sensor domain, leading to the compaction of the SpeFL effector domain and blocking the action of release factor RF1. Thus, our study not only reveals basic strategies by which nascent peptides assist the ribosome in detecting a specific metabolite, but also provides a framework for assessing how ornithine promotes virulence in several human pathogens.

## INTRODUCTION

Putrescine is a naturally abundant polyamine that is produced from ornithine by the enzyme ornithine decarboxylase, whose expression and activity are tightly regulated^2^. Two ornithine decarboxylase genes exist in *E. coli*, the constitutive *speC* and the inducible *speF*, which along with its operon partner *potE*, an ornithine-putrescine antiporter, is expressed under mild acidic stress and high ornithine levels^4–6^. Searching for regulatory elements upstream of *speF*, we found a short open reading frame (ORF) encoding a putative 34-amino acid peptide, which we named *speFL* (leader of *speF*) (see Methods, Fig. 1a). This ORF is conserved in many pathogenic γ-proteobacteria (Supplementary Fig. 1), including *Salmonella* Typhimurium (*S.* Typhimurium), where it was recently reported as *orf34*^7^. Translation of *orf34* in the presence of ornithine activates *speF* expression by preventing premature Rho-dependent transcription termination^7^. However, the mechanism by which ornithine triggers *speF* expression is unknown.

**Figure 1.**
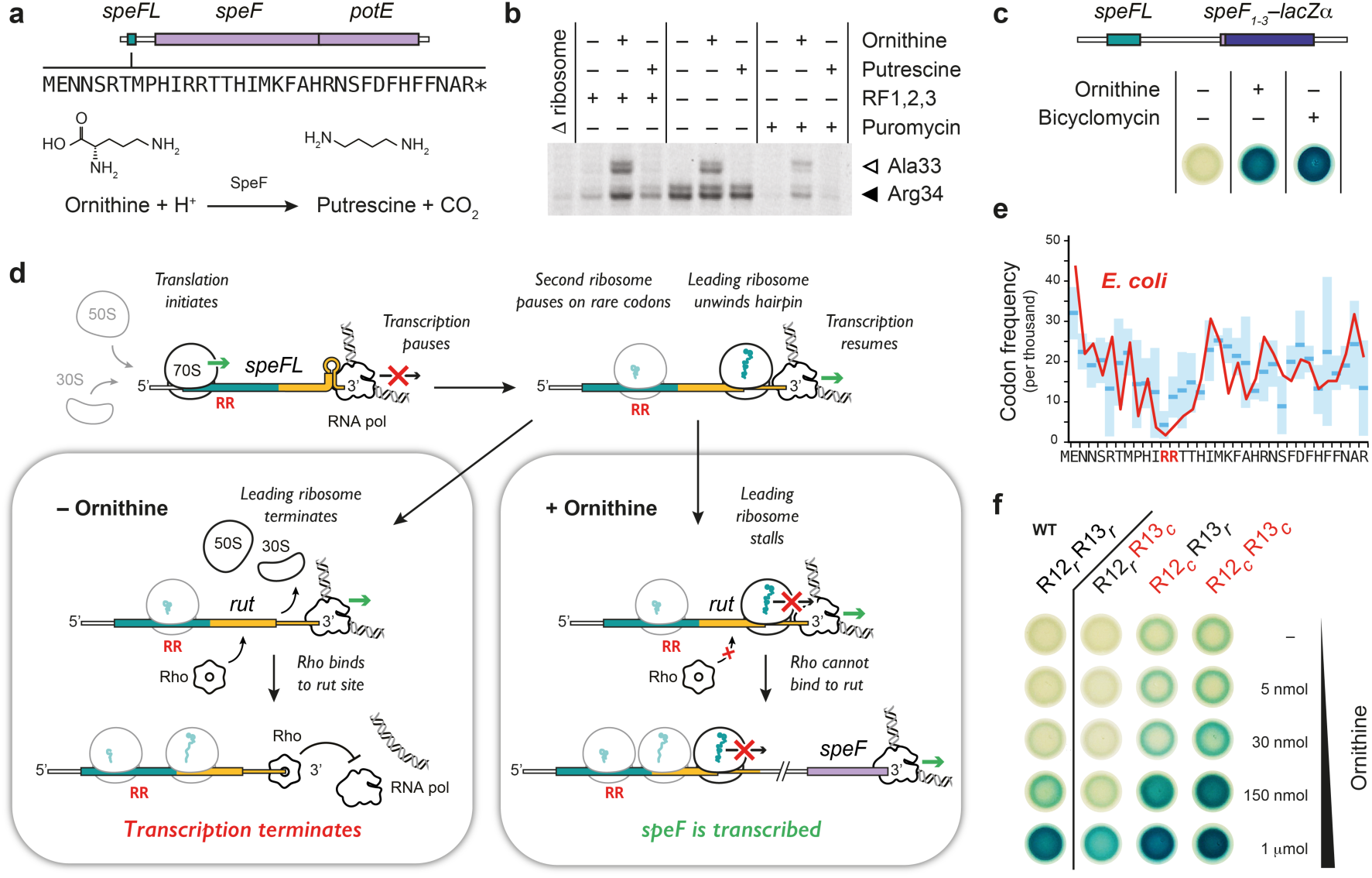
Mechanism of *speF* activation by SpeFL and ornithine. **a**, Schematic layout of the *speF* operon showing the sequence of the SpeFL peptide and the reaction catalyzed by SpeF. **b**, Toeprinting assay^8, 9^ to monitor the translation of *speFL* in the absence (–) or presence (+) of 10 mM ornithine, 10 mM putrescine, release factors (RF1,2,3) or 90 μM puromycin. Arrows indicate ribosomes stalled with the codon for the indicated amino acid in the P-site (Ala33 – open triangle; Arg34 – filled triangle). **c**, *E. coli* TB1 cells^56^ transformed with a plasmid carrying a *speF_1–3_-lacZα* translational fusion whose expression is placed under the control of *speFL* and the *speFL-speF* intergenic region. Cells were grown on rich medium supplemented with 50 µg/ml streptomycin, 100 µg/ml ampicillin, 1 mM IPTG and 0.5 mM X-Gal in the absence (–) or presence (+) of 3 μmol ornithine or 20 µg bicyclomycin. Blue cells express the *speF_1–3_-lacZα* translational fusion. **d**, Model of *speF* induction following the ornithine-dependent stalling of ribosomes translating *speFL*. The *speFL* open reading frame is boxed and shown partly in turquoise, with the overlapping *rut* site in yellow. Consecutive rare arginine codons R12 and R13 are shown in red letters. The leading ribosome on *speFL* is outlined in black while the second and third ribosomes are outlined in gray. The SpeFL peptide is in turquoise. **e**, Codon frequency at each position of *speFL* in *E. coli* (red line) and in the Enterobacteriales order (*n_species_*=10 (see Supplementary Fig. 1); mean – blue lines; ± standard deviation (SD) – light blue boxes). Codon usage values were obtained from the Codon Usage Database (NCBI-Genbank Flat File Release 160.0 [June 15 2007])^57^. **f**, The same assay as in c, showing the induction of a *speF_1–3_-lacZα* translational fusion by wild-type (WT – R12*_r_*R13*_r_*) or synonymous *speFL* variants with different combinations of rare (*r*) or common (*c*) arginine codons at positions 12 and 13, in the absence (–) or presence (+) of different amounts of ornithine. Mutated codons are shown in red.

## RESULTS

To investigate how *speF* is activated, we performed toeprinting assays^8, 9^ to monitor the position of ribosomes on a transcript encoding SpeFL (Fig. 1b and Supplementary Data 1, Supplementary Tables 1 and 2). A faint toeprint corresponding to ribosomes that reached the UAG stop codon was visible in the absence of exogenous ligand. Addition of ornithine resulted in two strong toeprint signals for ribosomes stalled with codons 33 or 34 of *speFL* in the ribosomal P-site. Ribosome stalling occurred in a dose-dependent manner with respect to ornithine concentration (Supplementary Fig. 2), but no toeprints were observed in the presence of putrescine, highlighting the strict dependence of the stalling process on ornithine availability. Translating a double-frameshifted *speFL_fs_* template that encodes a different amino acid sequence did not yield ornithine-dependent toeprints (Supplementary Fig. 3), indicating that ribosome stalling depends on the nascent peptide rather than on the mRNA structure. In the absence of release factors, the toeprint at position 34 intensified, reflecting impaired translation termination. Treatment with puromycin led to the disappearance of this toeprint, while ornithine-dependent toeprints remained visible. Puromycin causes premature peptide release and insensitivity to this antibiotic is characteristic of arrest peptides, a class of nascent regulatory peptides that stall the ribosomes that are translating them, often in a ligand-dependent manner^10, 11^. Finally, we showed that an RNA element including *speFL* and the 257-nucleotide *speFL-speF* intergenic region induces the expression of a *speF_1–3_-lacZα* translational fusion *in vivo* in response to ornithine (Fig. 1c). Treatment with bicyclomycin, which specifically blocks the ATPase activity of Rho^12^, resulted in constitutive *speF_1–3_-lacZα* expression, confirming the previously reported^7^ involvement of Rho in the regulation of *speF*. Thus, ribosomes translating *speFL* stall in an ornithine-dependent manner, inducing *speF* through a Rho-dependent mechanism.

In *S.* Typhimurium, Rho-dependent transcription termination occurs immediately downstream of an mRNA hairpin that includes the 3’ end of *speFL*^7^. This hairpin is conserved in *E. coli* and would cause an RNA polymerase that has just finished transcribing *speFL* to pause (Fig. 1d, Supplementary Fig. 4). A ribosome translating *speFL* can unwind the pause hairpin upon reaching the stop codon, freeing the RNA polymerase and allowing transcription to resume. When ornithine levels are low, the leading ribosome on *speFL* terminates and dissociates from the mRNA, exposing part of a predicted Rho utilization (*rut*) site^13^. Rho-mediated transcriptional attenuation is commonly used for metabolic control and can be coupled with translational arrest at a leader ORF, for example during the regulation of *E. coli* tryptophanase operon by the tryptophan-dependent arrest peptide TnaC^14^. For Rho to bind to the nascent transcript and cause premature transcription termination, a full *rut* site must be available. Since polysome accumulation on *speFL* would interfere with *rut* availability, we hypothesized that consecutive rare arginine codons at positions 12 and 13 of *speFL* may slow translation enough to fully expose the *rut* site and give Rho a chance to bind. As reported previously^7^, this region of *speFL* contains rare codons in many γ-proteobacteria, especially at position 12 (Fig. 1e). While replacing codon 13 with a common synonymous codon caused a mild decrease in *speF_1–3_-lacZα* induction, the same mutation at position 12 or mutation of both codons gave rise to a basal level of *speF* expression that was not observed with wild-type *speFL* (Fig. 1f), consistent with a model whereby efficient Rho binding is dependent on polysomes not accumulating on *rut*. When ornithine levels are high, the leading ribosome on *speFL* undergoes nascent peptide-mediated translational arrest. Ribosome stalling masks the *rut* site and polysomes accumulate (Supplementary Fig. 5). This prevents Rho from binding, allowing transcription to proceed and *speF* to be expressed.

To determine how nascent SpeFL functions as an ornithine sensor, we used cryo-EM to obtain two structures of a SpeFL–70S ribosome complex stalled in the presence of ornithine at an overall resolution of 2.7 Å (Fig. 2a, Supplementary Fig. 5, 6 and 7, and Supplementary Table 3). We observed a major subpopulation corresponding to a stalled termination complex with well-resolved density for a 34-residue peptidyl-tRNA^Arg^ bound to the P-site, but the elongation complex stalled on codon 33 that was seen by toeprinting was not observed (Fig. 2b). This discrepancy may result from differences in the kinetics of the purified translation system used for toeprinting and the S30 extract-based translation system used for cryo-EM sample preparation. Alternatively, the elongation complex may be less stable than the termination complex, resulting in complex dissociation during sucrose gradient centrifugation.

**Figure 2.**
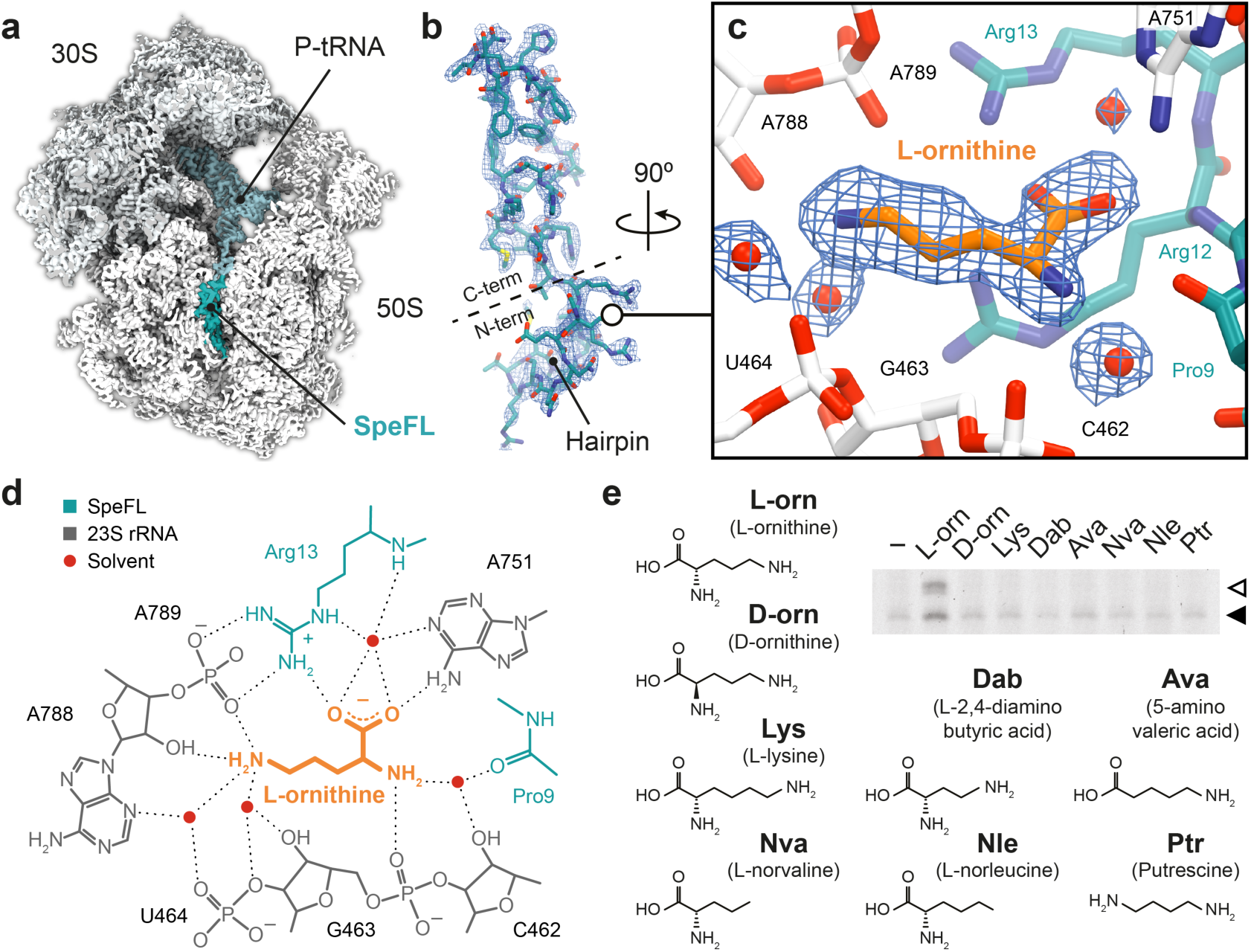
Structural basis for the specific recognition of L-ornithine by the SpeFL-70S complex. **a**, Transverse section of a cryo-EM density map of the SpeFL–70S complex, showing the small (30S, light gray) and large (50S, white) ribosomal subunits, the P-site tRNA (pale blue) and the SpeFL peptide (turquoise). **b**, Cryo-EM density displayed as a mesh, fitted with a molecular model of SpeFL, with the N– and C–terminal domains highlighted. The N-terminal hairpin is also indicated. **c**, Binding pocket formed by the 23S rRNA (white) and SpeFL (turquoise), with a single L-ornithine molecule (orange) surrounded by 4 solvent molecules (red) fitted into the cryo-EM density of the SpeFL-ESRF complex. The existence of these solvent molecules was validated using two independently determined structures of the SpeFL-70S complex (see Supplementary Fig. 9). **d**, Chemical diagram showing interactions between the 23S rRNA (dark gray), SpeFL (turquoise), L-ornithine (orange) and solvent molecules (red) inside the ligand binding pocket. Possible hydrogen bonds are shown as dotted lines. **e**, Toeprinting assay^8, 9^ to monitor the translation of *speFL* in the absence (–) or presence of 10 mM (+) of various small molecules in the presence of release factors (RF1,2,3). Arrows indicate ribosomes stalled with the indicated amino acid in the P-site (Ala33 – open triangle; Arg34 – filled triangle).

In the stalled SpeFL-70S termination complex, SpeFL adopts a compact fold that completely obstructs the upper two-thirds of the exit tunnel and can be subdivided into N-and C-terminal domains, corresponding to residues 1–13 and 14–34 of SpeFL, respectively. The N-terminal domain forms a hairpin, while secondary structure elements stabilize the C-terminal domain, most notably two type I β-turns between residues 19–22 and 23–26, and one 3_10_–helix between residues 27–32. In addition, SpeFL interacts extensively with the 23S ribosomal RNA (23S rRNA) and with ribosomal proteins uL4 and uL22 through a combination of π–stacking, salt bridges and hydrogen bonding (Supplementary Fig. 8). All of these structural elements contribute to stabilizing the complex fold adopted by SpeFL inside the exit tunnel.

A clear peak of density that could be unambiguously attributed to a single L-ornithine molecule was visible inside a cavity formed by 23S rRNA residues C462, G463, U464, A751, A788 and A789 and by Pro9, Arg12 and Arg13 of the N-terminal domain of SpeFL, referred to here as the sensor domain (Fig. 2c, d and Supplementary Fig. 9). To our knowledge, this cavity represents a novel binding site for small molecules on the ribosome, which could be targeted for future antibiotic development (Supplementary Fig. 10). The ornithine recognition loop of SpeFL consists of a HIRRXXH motif spanning residues 10–16, among which His10, Arg13 and His16 help form the ligand-binding pocket by interacting with 23S rRNA and ribosomal protein uL22 residues (Fig 2c, d and Supplementary Fig. 8). Deletion of residues 1–7 of SpeFL, which disrupts the hairpin but retains the HIRRXXH motif (Supplementary Fig. 11), mutation of the strictly conserved Arg12 and Arg13 to alanine or lysine (Supplementary Fig. 12) or mutation of His10, Ile11 or His16 to alanine (Supplementary Fig. 13) abolished ornithine-dependent translational arrest *in vitro*, highlighting the importance of the hairpin and of the conserved residues of the HIRRXXH motif for the stalling process. The side chain and α-amino groups of ornithine interact with the backbone phosphates of 23S rRNA residues A789 and G463, respectively (Fig. 2d). Ornithine is further stabilized via hydrogen bonding between its α-carboxyl group and both the guanidino group of SpeFL residue Arg13 and the N6-amino group of 23S rRNA residue A751. The availability of two independently obtained cryo-EM maps of the SpeFL-70S complex enabled us to accurately model and cross-validate the positions of ordered metal ions and solvent molecules within the complex. Four of these solvent molecules fill the cavity and make additional bridging interactions between ornithine and the 23S rRNA or SpeFL (Fig. 2c, d and Supplementary Fig. 9). Thus, small molecules differing from ornithine by only a single methylene group are either too short (L-2,4–diaminobutyric acid) or too long (L-lysine) for the binding pocket, while the deletion of ligand functional groups (putrescine, L-norvaline, L-norleucine, 5-aminovaleric acid) or the use of a D-enantiomer (D-ornithine) abolish stalling by preventing the formation of certain hydrogen bonds (Fig. 2e and Supplementary Data 2). The tight coordination of L-ornithine via each of its potential hydrogen bond donors and acceptors therefore explains the high selectivity of SpeFL and the ribosome for their cognate ligand.

To understand how ornithine capture by the sensor domain stalls the ribosome, we must focus on the C-terminal effector domain of SpeFL, which consists of a hydrophobic core composed of four phenylalanine residues (Phe20, Phe28, Phe30 and Phe31) nucleated around the strictly conserved Phe26 (Fig. 3a and Supplementary Fig. 8). Residues Phe28, Phe30 and Phe31 establish π–stacking interactions with the bases of 23S rRNA residues U2586, G2505 and A2062, respectively, which help to position the effector domain in the upper part of the ribosomal exit tunnel (Fig. 3a, Supplementary Fig. 8). Mutation of Phe26 or any of these three aromatic residues to alanine abolishes or severely impairs ribosome stalling *in vitro* (Fig. 3b and Supplementary Data 3), highlighting their importance for translational arrest. Since the SpeFL-70S structure corresponds to stalled ribosomes with a UAG stop codon in the A-site, it is clear that SpeFL must inhibit the action of RF1, the release factor responsible for recognizing this stop codon. Comparing our structure with that of a *Thermus thermophilus* 70S ribosome in complex with RF1 and a P-site tRNA^15^ reveals that the binding of RF1 to the SpeFL-70S complex is prevented by 23S rRNA residue U2585, which adopts a conformation that would sterically clash with the GGQ loop of RF1. The movement of U2585 is caused by residue Asn32 of SpeFL, which takes the place of its base in the 70S–RF1–P-tRNA complex (Fig. 3c, d). The nature of residue 32 is likely not to be critical for stalling as even an N32A mutant undergoes ornithine-dependent translational arrest (Fig. 3b and Supplementary Data 3). Moreover, the identity of the release factor is not important as all possible stop codons are observed for *speFL* across different species (Supplementary Fig. 1). Thus, the continued synthesis of SpeFL after the recognition of ornithine by the sensor domain leads to the compaction of the effector domain and forces U2585 into a conformation that prevents release factor action, causing the ribosome to stall.

**Figure 3.**
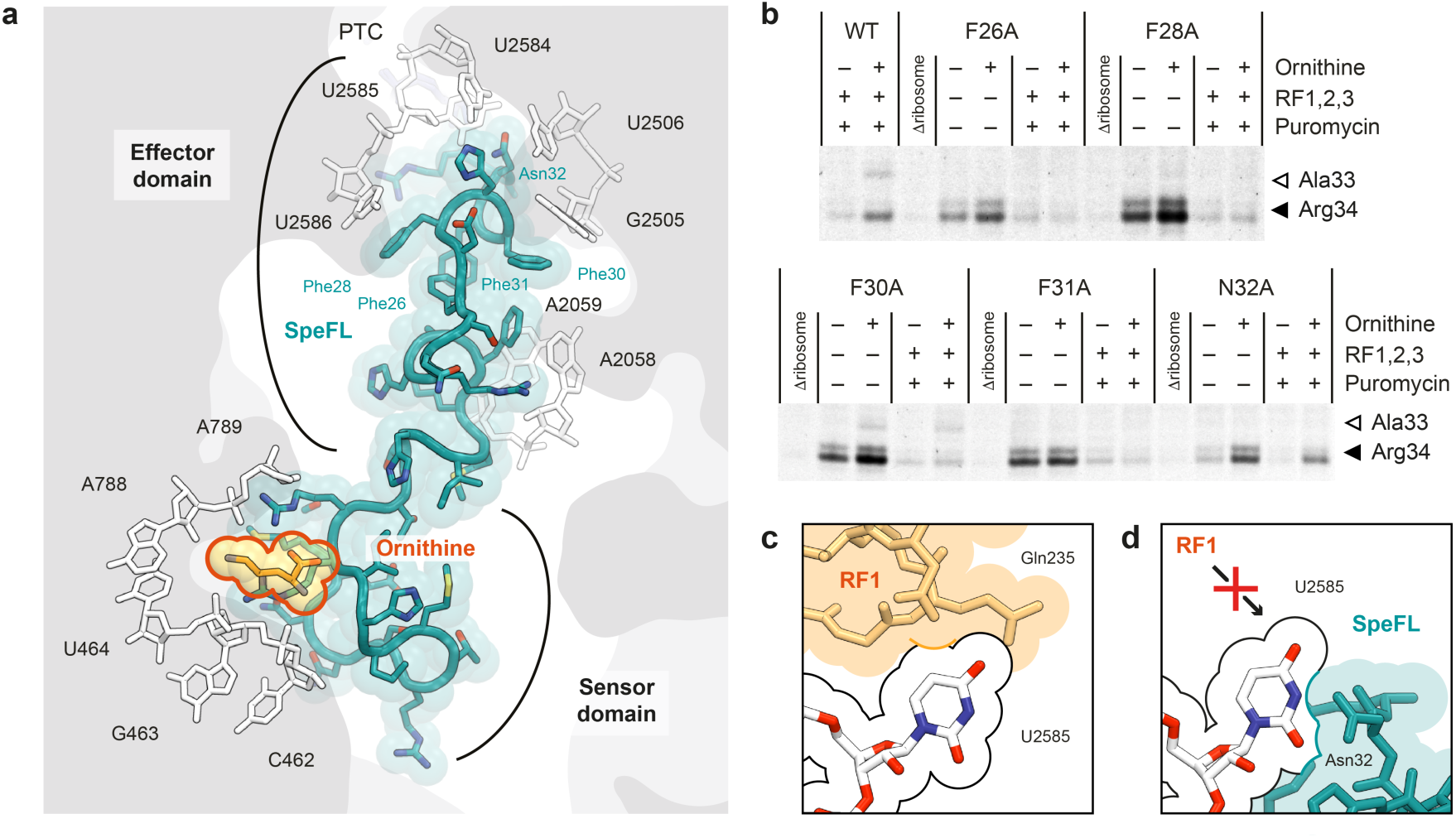
Inhibition of peptide release by SpeFL. **a**, Close-up of the ribosomal exit tunnel showing the sensor and effector domains of SpeFL (turquoise) interacting with residues of the 23S rRNA (white). Ornithine (orange) is trapped between the tunnel wall and the sensor domain, while synthesis of all 34 amino acids of SpeFL leads to compaction of the effector domain and blockage of the peptidyl transferase center (PTC). **b**, Toeprinting assay^8, 9^ to monitor the translation of wild-type (WT) and mutant *speFL* in the absence (–) or presence (+) of 10 mM ornithine or 90 µM puromycin. Arrows indicate ribosomes stalled with the indicated amino acid in the P-site (Ala33 – open triangle; Arg34 – filled triangle). **c**, Structure of an *E. coli* 70S–RF1–P-tRNA complex (PDB 5J3C)^15^, showing the GGQ loop of RF1 (peach, with residue Gln-235 labeled) and 23S rRNA residue U2585 (white). **d**, Close-up of the SpeFL–70S structure showing the same view as in c. Residue Asn32 of SpeFL (turquoise) forces U2585 to adopt a conformation that prevents RF1 binding.

Metabolite sensing by a translating ribosome is a complex and dynamic process whose understanding has been hampered by the lack of high-resolution structural data^11, 16–18^. With the exception of antibiotic-dependent translational arrest^19–21^, in which the drug binds directly to the empty ribosome, it is not known if the metabolite helps to create a binding surface for the nascent peptide or vice versa^11^, even though the latter has been suggested for the binding of tryptophan to the nascent TnaC peptide^17^. In the SpeFL–70S structure, ornithine interacts primarily with the ribosome, either directly or via bridging solvent molecules, whereas SpeFL provides only a few stabilizing interactions that help capture the cognate ligand (Fig. 2d). This implies that ornithine is already loosely associated with the 23S rRNA prior to the arrival of the SpeFL sensor domain (Fig. 4), as suggested by molecular dynamics simulations pointing to the existence of binding crevices for different amino acid side chains within the ribosomal exit tunnel^22^. Additionally, the close proximity of the sensor domain to the tunnel constriction formed by ribosomal proteins uL4 and uL22 raises the possibility that the sensor domain begins to fold in the upper part of the exit tunnel, consistent with the decreased *speF* expression seen for the R12_r_R13_c_ mutant (Fig. 1f). Indeed, a partially folded sensor hairpin emerging from the tunnel constriction could rapidly contact and fix an ornithine molecule present within its adjacent binding crevice on the 23S rRNA. Once the interaction between the sensor domain and the tunnel wall has been stabilized through the binding of ornithine, the effector domain can be synthesized and compacted, resulting in the inhibition of peptide release from the ribosome. Since metabolite recognition must occur before SpeFL is fully synthesized, this process has to operate under a kinetic regime. In other words, the concentration of free ornithine must be significantly above the K_D_ of the interaction, resulting in a rate of metabolite binding that is greater than the rate of SpeFL synthesis. This increases the likelihood that ligand recognition takes place by ensuring that ornithine is present within its binding pocket when the sensor domain of SpeFL reaches the corresponding region of the exit tunnel. Such a scenario is reminiscent of kinetically controlled riboswitches, which require high ligand concentrations to ensure that metabolite binding occurs faster than RNA transcription^23^. These basic strategies for ligand recognition could be used by other metabolite-sensing nascent peptides.

**Figure 4.**
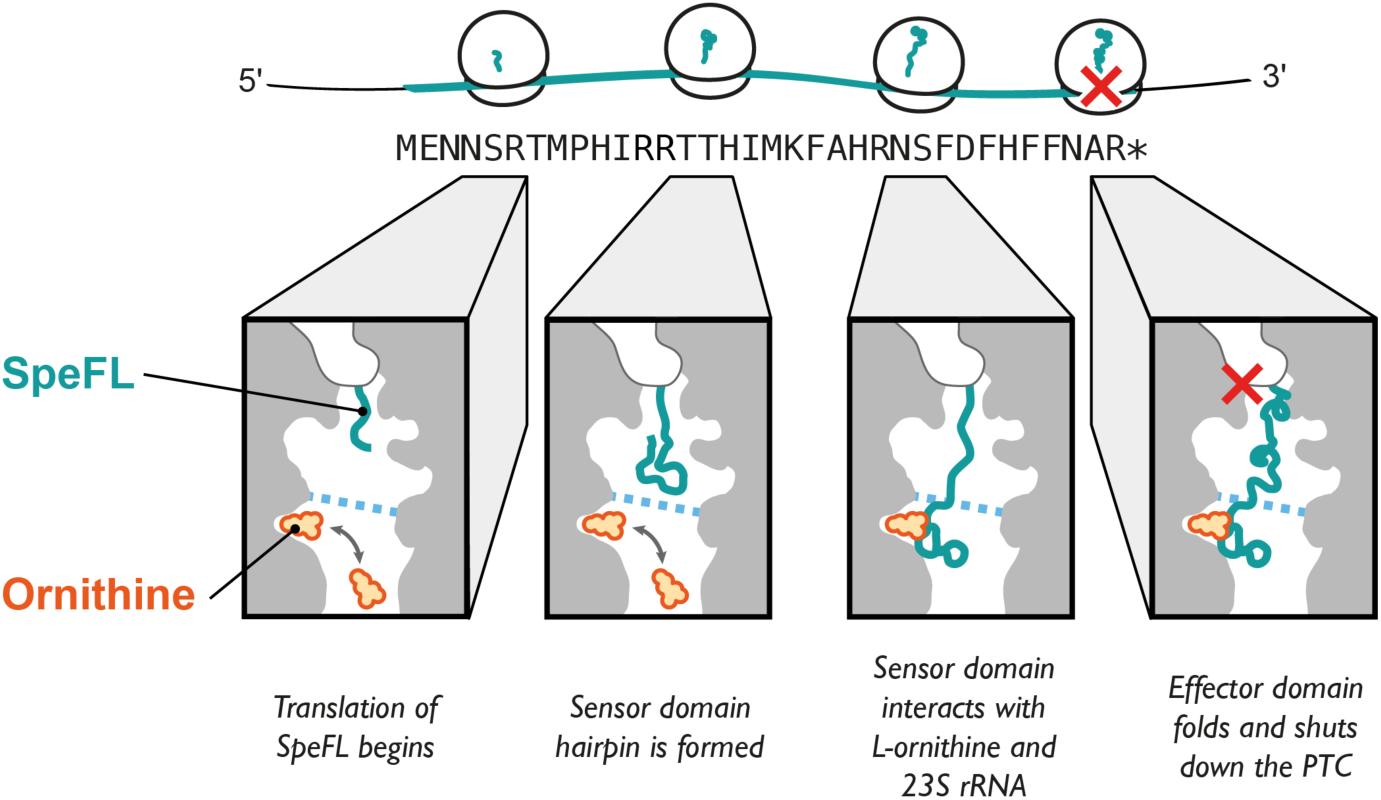
Mechanism of ornithine sensing and capture by the SpeFL–70S complex. Model for the binding of ornithine (orange) by the ribosome and SpeFL (turquoise), leading to inactivation of the peptide release activity of the ribosome (red cross). The tunnel constriction is shown as a blue dotted line.

In conclusion, our data reveal how the ribosome aided by SpeFL functions as a highly selective ornithine sensor to regulate polyamine biosynthesis in pathogenic bacteria like *E. coli* or *S.* Typhimurium. Both *speF* and *potE*, whose expression is regulated by SpeFL in response to fluctuating ornithine levels, have been linked to biofilm formation and microbial virulence^24–26^. However, the exact physiological roles of the *speF* operon remain unclear. A recent study on interspecies niche modulation during mixed *Enterococcus faecalis* (*E. faecalis*) and uropathogenic *E. coli* infections^27^ may provide us with a clue in this regard. In this work, ornithine secreted by *E. faecalis* helps uropathogenic *E. coli* grow in the iron-restricted environment of a human host by redirecting its metabolism towards siderophore production. Since we know that *E. faecalis* lowers the pH of the polymicrobial culture by producing lactic acid and that low pH and ornithine are required for *speF* induction, it is reasonable to assume that SpeFL could play a role in this *E. faecalis*-driven promotion of *E. coli* growth. Indeed, ornithine produced by *E. faecalis* might be sensed by SpeFL to trigger the expression of SpeF, which in turn decarboxylates ornithine into putrescine. Putrescine turns on the expression of a number of genes^28^, including the sigma factor FecI, a protein that activates the ferric citrate uptake operon *fecABCDE*. Ferric citrate is a naturally abundant compound inside the host, which allows uropathogenic *E. coli* to grow under iron limiting conditions^29^. Although a direct link between ornithine secreted by *E. faecalis* and ferric citrate uptake has yet to be demonstrated, it is known that the production of the siderophores enterobactin and yersiniabactin is stimulated in this manner^27^. Thus, a possible involvement of SpeFL in sensing ornithine cues within polymicrobial communities of this kind must now be examined.

## Acknowledgement

A.H.V. is funded by a doctoral grant from the French Ministère de l’Enseignement Supérieur et de la Recherche. C.A.I., B.S. and G.S. have received funding for this project from the European Research Council (ERC) under the European Union’s Horizon 2020 research and innovation program (Grant Agreement No. 724040). C.A.I. is an EMBO YIP and has received funding from the Fondation Bettencourt-Schueller. I.C-M. is funded by Inserm and A.C.S. is funded by a joint doctoral grant from Inserm and the Aquitaine Regional Council. We acknowledge Diamond for access and support of the Cryo-EM facilities at the UK national electron bio-imaging centre (eBIC), (proposal EM 19716-1), funded by the Wellcome Trust, MRC and BBSRC, and Sarah Neumann, Andrew Howe and James Gilchrist for assistance. We also acknowledge the European Synchrotron Radiation Facility for provision of microscope time on CM01^30^ and we thank Michael Hons for his assistance. This work has been supported by *iNEXT*, grant number 3901, funded by the Horizon 2020 program of the European Union. We thank Isabelle Iost for help with gradient fractionation, Yaser Hashem, Armel Bezault and Marion Decossas for help with grid preparation and screening, Rémi Fronzes for access to the GPU cluster, Chiara Rapisarda for help with cryo-EM data processing and Nora Vazquez-Laslop for providing the *E. coli* TB1 strain and pErmZα plasmid for the *in vivo* lacZ assay.

## Author contribution

C.A.I. and B.S. designed the study. I.C-M. identified *speFL*. A.H.V., B.S., G.S. and A.C.S. performed biochemical experiments. A.H.V. performed bacterial assays. A.H.V. prepared the cryo-EM sample. A.H.V. and C.A.I. processed the cryo-EM data. A.H.V., B.S., G.S., A.C.S. and C.A.I. interpreted the results. A.H.V., B.S. and C.A.I. wrote the paper.

## METHODS

No statistical methods were used to predetermine sample size. The experiments were not randomized and investigators were not blinded to allocation during experiments and outcome assessment.

### Bioinformatic identification of *speFL*

Homologs of *E. coli speF* were identified using *tblastn*^31^ (E-value lower than 10^-4^, >70% coverage), redundancy was minimized using *CD-Hit*^32^ (95% sequence identity cutoff) and regions between *speF* and the nearest upstream annotated genes were compiled (unknown, hypothetical, uncharacterized or leader genes were considered not to be annotated). All possible forward ORFs within the last 500 nucleotides of these upstream regions were extracted, considering ATG and alternative start codons defined for bacterial, archaeal and plant plastid genetic codes (NCBI Genetic codes Table 11). Possible Shine-Dalgarno sequences were not taken into account. In cases where more than one ORF was possible, the longest ORF was kept. Redundancy within the ORFs thus obtained was minimized with *CD-Hit* (95% sequence identity cutoff). A pairwise comparison of the resulting ORFs was then carried out as follows. First, nucleotide sequences were translated into amino acid sequences. A sliding window of 10 amino acids was applied to each sequence and an alignment score based on a BLOSUM62^33^ substitution matrix was computed for all possible combinations of 10-amino acid fragments from each pair of ORFs (no gaps allowed). A graph in which each node represents a 10-amino acid fragment and each edge represents an alignment score greater than 10 between two fragments (with the alignment score as a weight) was constructed. Finally, we used *MCL*^34, 35^ to identify clusters within this graph and found a major cluster of conserved upstream ORFs corresponding to *speFL*.

### Phylogeny of *speFL*

Homologs of *E. coli speFL* were identified using *tblastn*^31^ and a phylogenetic analysis of *speFL* from 15 representative species of γ–proteobacteria was carried out using the EMBL-EBI *Simple Phylogeny* server^36^. The resulting tree was displayed with *Dendroscope*^37^ (Supplementary Fig. 1).

### Toeprinting assays

Toeprinting was performed as described previously^9^. Briefly, DNA templates containing a T7 promoter, a ribosome binding site, wild-type or mutant *speFL*, the first 75 nucleotides of the *speFL-speF* intergenic region and the NV1 sequence^38^ were generated or amplified by polymerase chain reaction (PCR). The wild-type template was amplified by PCR from *E. coli* DH5α genomic DNA using oligonucleotides 1, 2 and 3 followed by PCR amplification of the product with oligonucleotides 4 and 5 (see Supplementary Table 1 for the sequences of all oligonucleotides used). The *speFL*-Δ1–7 template was obtained using the same protocol as for the wild type, but substituting oligonucleotide 2 with 6. Templates with point mutations and the double frame-shifted template *speFL_fs_* were purchased as gBlocks from IDT and amplified with oligonucleotides 4 and 5 (see Supplementary Table 2 for the sequences of all DNA templates). DNA templates were transcribed and translated *in vitro* using the PURExpress Δ RF123 system (New England Biolabs). Ligands were dissolved in water and added as needed at the beginning of the reaction. A Yakima Yellow-labeled probe (2 µM) complementary to the NV1 sequence^38^ was added to the 5 µL reaction after incubating for 60 minutes at 30°C, and the sample was incubated for another 5 minutes at the same temperature. When needed, samples were treated with 90 µM puromycin at 30°C for 3 minutes, immediately followed by reverse transcription with 50 U of AMV reverse transcriptase (Promega) for 20 minutes at 30°C. RNA was degraded by adding 0.5 µL of a 10 M NaOH stock at 30°C for 15 minutes. Samples were neutralized with 0.7 µL of a 7.5 M HCl stock and the remaining cDNA was purified using a nucleotide removal kit (Qiagen). Sequencing reactions were performed according to the method of Sanger. Briefly, 1 pmol of DNA template was mixed with 10 pmol of oligonucleotide 9 labeled with Yakima Yellow and 1µL of HemoKlen Taq DNA Polymerase (New England Biolabs) in a 6µL reaction mixture containing 50 mM Tris-HCl pH 9.0, 2 mM MgCl_2_, 6.6 µM dNTPs, 10 µM ddGTP, 117 µM ddATP, 200 µM ddTTP or 66 µM ddCTP. Primers were extended with 30 cycles of 30 seconds of annealing at 42 °C and 1 minute of elongation at 70 °C. The purified cDNA and sequencing reactions were dried using a SpeedVac and resuspended in 6 µl or 3.5 µl gel-loading buffer (95 % formamide, 0.25 % (w/v) xylene cyanol, 0.25 % (w/v) SDS), respectively. Samples were denatured at 95°C for 5 minutes, and 2µL of the sequencing reactions and 3 µL of the toeprinting were separated by 7.5 % sequencing PAGE (2000 V, 40 W for 2-2.5h) followed by detection on a Typhoon imager (GE).

### β-galactosidase assay

To test for *in vivo* activity, a translational reporter plasmid was obtained by fusing a region containing *speFL*, the *speFL–speF* intergenic region and the first three codons of *speF* to lacZα. The insert was prepared by PCR amplification from the *E. coli* K12 genome using oligonucleotides 7 and 8. Oligonucleotides 9 and 10 were used to linearize the pErmZα plasmid^39^. The insert and linearized plasmid were mixed and transformed following the AQUA cloning protocol^40^. Plasmids containing point mutations in the R12–R13 region were generated by site-directed mutagenesis as follows. The wild-type plasmid was linearized by PCR amplification with oligonucleotides 11 and 12, 13 or 14 (the latter included the mutations). The PCR product was purified from a 2% TAE-agarose gel with a Gel Extraction Kit (Qiagen) and phosphorylated for 30 minutes at 37 °C with 4U of T4 Polynucleotide Kinase (New England Biolabs) in a total volume of 20 µl according to the manufacturer’s instructions. The plasmid was circularized again by incubating the phosphorylated product with 400 U of T4 DNA ligase (New England Biolabs) for 2 hours at 16°C. The plasmids were transformed into *E. coli* TB1^39^ and the cells were grown in lysogeny broth (LB) at 37°C (200 rpm) with streptomycin (50 µg/ml) and ampicillin (100 µg/ml) until they reached an optical density of 0.6 at 600 nm. 5 µL of the cell culture were plated onto LB-agar plates supplemented with streptomycin, ampicillin, 1 mM Isopropyl-β-D-1-thiogalactopyranoside (IPTG) and 0.5 mM 5-bromo-4-chloro-3-indolyl-beta-D-galactopyranoside (X-gal). 20 µg of bicyclomycin (Santa Cruz Biotechnology) or 0–3 µmol of L-ornithine (Sigma) were added after a 6 hour incubation at 37°C. The plates were then incubated at 37°C overnight and pictures were taken the next day.

### Preparation of an *E. coli* SpeFL–70S complex for cryo-EM

The SpeFL–70S complex was prepared using a modified disome purification strategy^20^. Briefly, SpeFL was expressed in an RTS 100 *E. coli* HY Kit (Biotechrabbit) for 1 hour at 30°C in the presence of 10 mM L-ornithine if indicated, using a pEX-K4-SpeFL_2x plasmid (Eurofins) that carries two copies of *speFL* arranged as a bicistronic mRNA (Supplementary Fig. 5; insert sequence: 5’-CGA-TCG-AAT-TCT-AAT-ACG-ACT-CAC-TAT-AGG-GCT-TAA-GTA-TAA-GGA-GGA-AAA-AAT-ATG-GAA-AAT-AAC-AGC-CGC-ACT-ATG-CCC-CAT-ATA-AGG-CGG-ACA-ACT-CAT-ATT-ATG-AAG-TTT-GCT-CAT-CGC-AAT-AGC-TTC-GAC-TTT-CAC-TTC-TTC-AAT-GCC-CGT-TAG-TCT-ACC-GAC-TAA-GGG-CAC-TTC-AGC-TAA-AGT-TTT-ATA-AGG-AGG-AAA-AAA-TAT-GGA-AAA-TAA-CAG-CCG-CAC-TAT-GCC-CCA-TAT-AAG-GCG-GAC-AAC-TCA-TAT-TAT-GAA-GTT-TGC-TCA-TCG-CAA-TAG-CTT-CGA-CTT-TCA-CTT-CTT-CAA-TGC-CCG-TTA-GTC-TAC-CGA-CTA-AGG-GCA-CTT-CAG-CTA-GAT-ATC-TAG-CAT-AAC-CCC-TTG-GGG-CCT-CTA-AAC-GGG-TCT-TGA-GGG-GTT-TTT-TG-3’). Reaction volumes of 50 µL and 750 µL were used for analytical and preparative purposes, respectively. When indicated, the reaction was treated with 100 μM puromycin for 3 minutes at 30°C before being layered over 10-40% (w/v) sucrose gradients containing Buffer A (50 mM HEPES-KOH pH 7.5, 100 mM K-acetate, 25 mM Mg-acetate and 10 mM L-ornithine), prepared using a Gradient Master 108 (Biocomp). Sucrose gradient ultracentrifugation was performed for 2 hours and 45 minutes at 35,000 rpm in a SW 41 Ti rotor (Beckman-Coulter) at 4°C. Polysome fractions were detected and collected using a UV detection system (UA-6, Teledyne ISCO) coupled to a gradient fractionator (Foxy R1, Teledyne ISCO). Polysomes were washed in 100 kDa molecular weight cutoff (MWCO) spin concentrators to remove sucrose, concentrate ribosomes and replace the solution with Buffer A. The concentration of ribosomes was inferred by measuring the absorbance of the sample at 260 nm (1 A_260_ = 60 μg/ml or 24 nM) with a NanoDrop One (ThermoFisher). For analytical purposes, 13.2 pmol of ribosomes with an excess of 10 nmol of *rnaseH* oligonucleotide were incubated for 1 hour at 25°C with 7.5 U of RNase H or without it (RNase H– control). The sample for cryo-EM grid preparation was treated with 75 U of RNase H (New England Biolabs) per 250 pmol of ribosomes for one hour at 25°C in the presence of 5 nmol of oligonucleotide 15. The monosomes obtained after RNAse H treatment were isolated by sucrose gradient ultracentrifugation as described above. The sample was flash frozen in liquid nitrogen and stored at −80 °C.

### Cryo-EM grid preparation

Frozen SpeFL–70S complex was thawed and diluted in Storage Buffer to yield a final concentration of 120 nM. Quantifoil carbon grids (QF-R2/2-Cu) were coated with a thin carbon layer prepared using an Edwards Vacuum Carbon Coater E306. Grids were glow discharged for 30 seconds at 2 mA before application of 4 μL of the SpeFL–70S complex. After blotting for 2 seconds and waiting for 30 seconds, grids were plunge-frozen in liquid ethane using a Vitrobot (FEI) set to 4°C and 100% humidity.

### Cryo-EM data acquisition and processing

Grids were imaged using two 300-keV Titan Krios (FEI) equipped with a K2 Summit direct electron detector (Gatan) at ESRF (France) and at the Diamond Light Source (eBIC, UK) producing the SpeFL-ESRF and SpeFL-DLS datasets, respectively. Images were recorded with EPU in counting mode with a magnified pixel size of 1.067 Å (Supplementary Table 3). 30 frames were collected to have a total accumulated dose of 30 electrons per Å^2^. Data were processed in *Relion 2.1*^41^, *Relion 3.0*^42^ and *Cryosparc 0.6*^43^ according to the scheme presented in Supplementary Figure 6. Briefly, MotionCor2^44^ was used for movie alignment, *Gctf*^45^ for Contrast Transfer Function (CTF) estimation and either *Relion 2.1* or *Cryosparc* for 2D classification of the particles obtained by automated picking in *Relion 2.1*. The *csparc2star.py* and *star.py* scripts^46^ were used for exporting particles selected in *Cryosparc* back into *Relion*. 3D classification was performed in *Relion 2.1* in three steps: (i) unsupervised classification with 4 times downsized particles, (ii) focused classification on all 3 tRNA sites with background subtraction and 3 times downsized particles, and (iii) focused classification on the P-site tRNA with background subtraction and 2 times downsized particles. Classes containing a single P-tRNA or both P-and E-site tRNAs were combined after ensuring that each class contained a peptide with the same conformation in the ribosomal exit tunnel when refined individually. Movie refinement and particle polishing were first performed with *Relion 2.1*. Refined particle coordinates were then used to re-extract particles in *Relion 3.0* in order to perform per particle CTF and beam tilt refinement, followed by Bayesian polishing.

### Model building and refinement

An initial model of the SpeFL–70S complex was obtained by placing the coordinates for an *E. coli* 70S ribosome (PDB 4U27)^47^ into the cryo-EM density map with *Situs*^48^, using the *colores* routine for the initial fit at 15 Å and the *collage* routine for fitting subdomains of the ribosome (30S body, 30S head, 30S spur, 50S body and L1 stalk) as independent rigid bodies at progressively higher resolutions until reaching the map resolution. The pixel size was optimized by generating post-processed maps with different pixel sizes in *Relion 2.1* and assessing the map-to-model correlation after real space refinement in *Phenix*^49^ with the initial model. A model for the SpeFL peptide was built manually with *Coot*^50^ and refined through multiple rounds of real-space refinement in *Phenix* and manual rebuilding in *Coot*. The model was validated with *MolProbity*^51^. Automatic map sharpening was performed in *Phenix* using a refined model from which L-ornithine had been removed (Supplementary Fig. 7 and 9). The quality of the sharpened maps was such that ordered ions and solvent molecules could be modeled into the density and validated using the two structural replicates. Hydrated magnesium ion clusters were identified on the basis of their distinctive cryo-EM density. The identity of individual magnesium and potassium ions could be inferred by analogy with a recent X-ray structure of the *Thermus thermophilus* ribosome^52^. All other solvent molecules were modeled as unknown atom and residue types. The resulting map was used to prepare all figures except Fig. 2a, for which a post-processed map from *Relion* was used (sharpening B-factor of –10).

### Figure preparation

Figures showing cryo-EM density or atomic models were prepared using *Chimera*^53^, *Chimera X*^54^ or *Pymol Molecular Graphics Systems* (version 1.7.4 Schrödinger)^55^.

### Data availability

The SpeFL-ESRF and SpeFL-DLS structures were deposited in the RCSB PDB with accession codes 6TC3 and 6TBV, and cryo-EM maps were deposited in the EMDB with accession codes EMD-10458 and EMD-10453.

## SUPPLEMENTARY INFORMATION

**Supplementary Figure 1.**
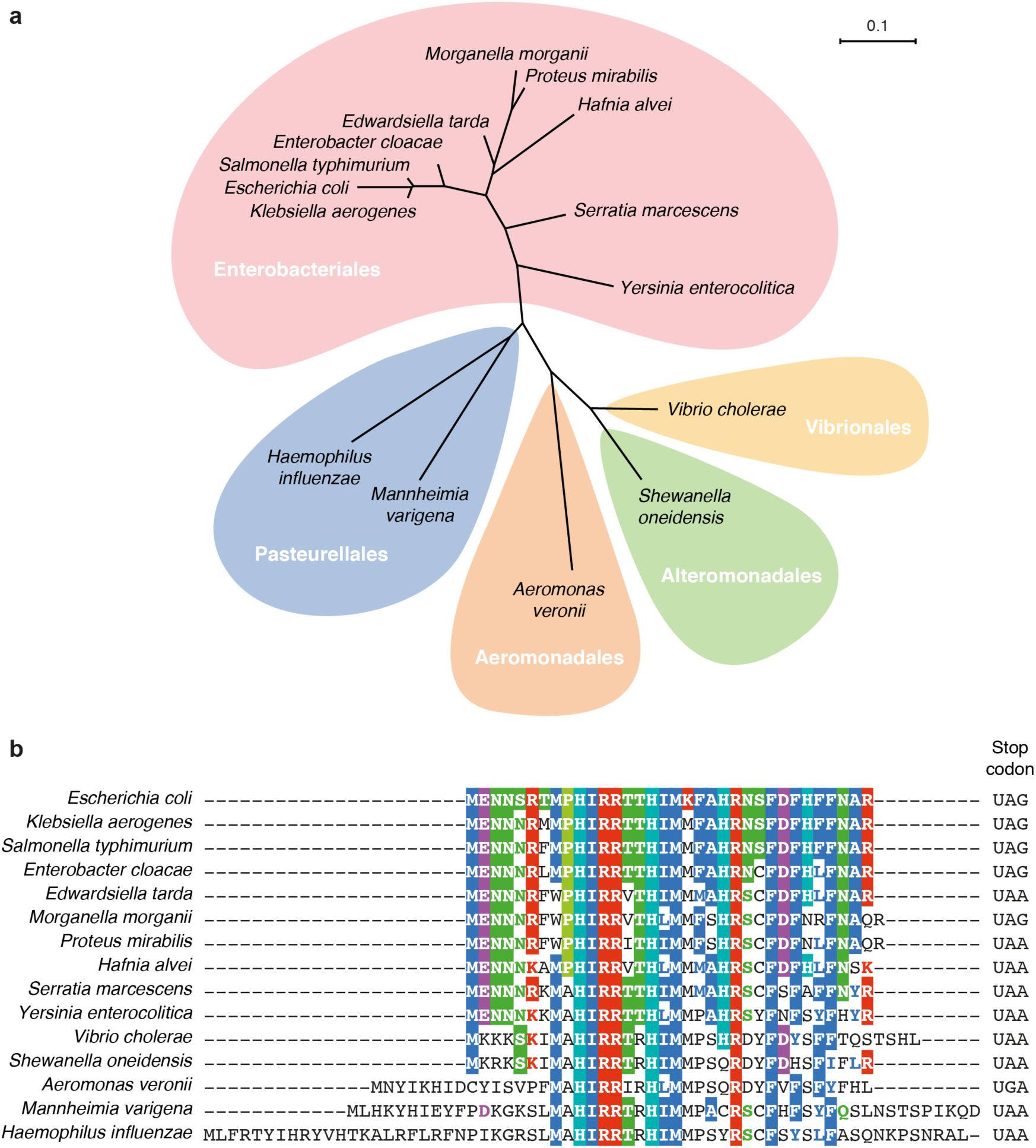
Conservation of SpeFL across γ-proteobacteria. **a**, Phylogenetic tree showing the distribution of representative *speFL* sequences from several orders of γ-proteobacteria. **b**, Multiple sequence alignment of SpeFL homologs from different species. The sequence shown for *Salmonella typhimurium* corresponds to the previously reported *orf34*^1^. SpeFL and its homologs belong to the group of proteins of unknown function DUF2618^2^. The stop codon found after each sequence is indicated on the right of the alignment.

**Supplementary Figure 2.**
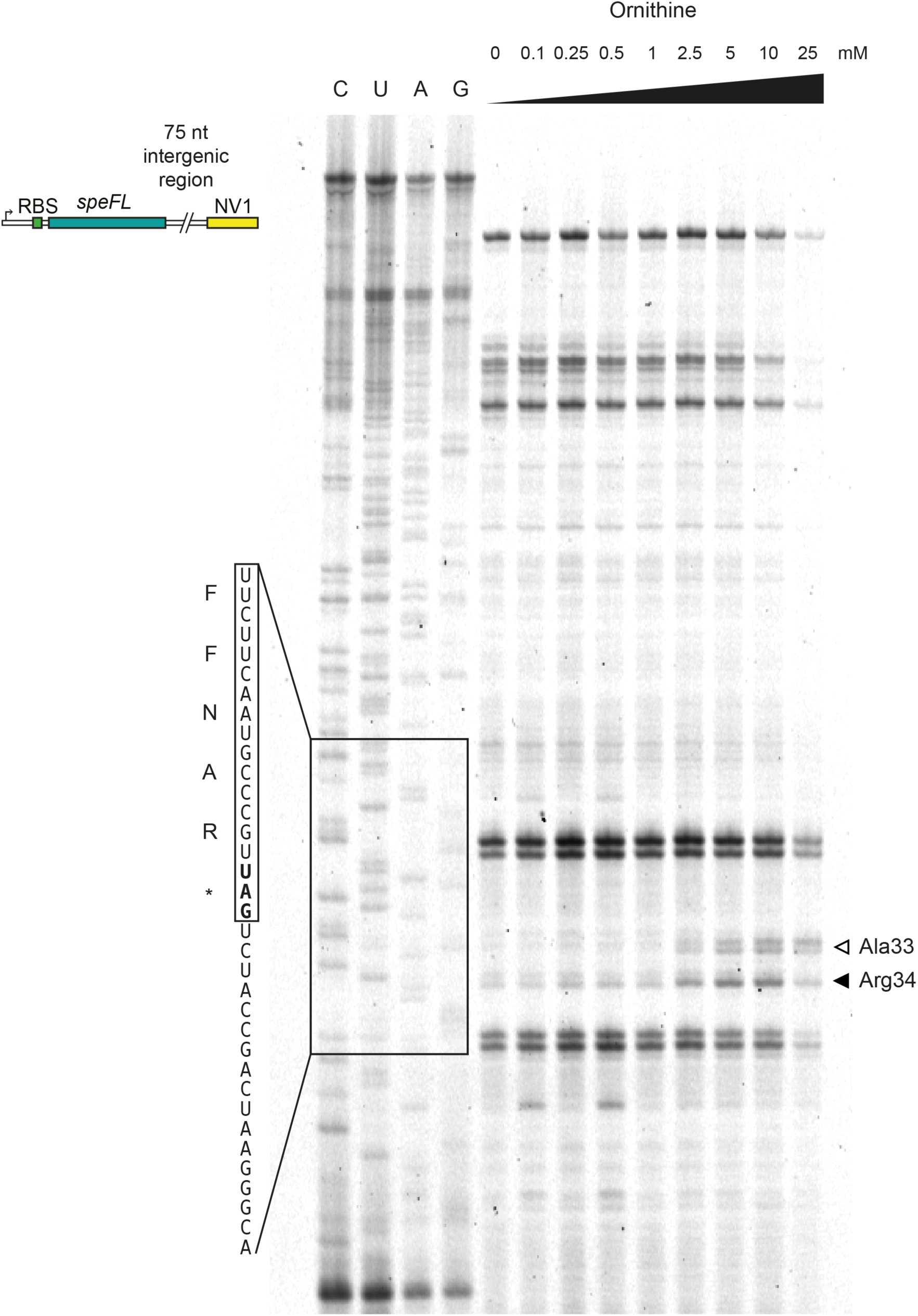
Dose-dependence of ornithine-mediated ribosome stalling on *speFL*. Toeprinting assay^3, 4^ to monitor the translation of wild type *speFL* in the presence of increasing concentrations of ornithine. All samples were treated with 90 µM puromycin. Arrows indicate ribosomes stalled with the codon for the indicated amino acid in the P-site (Ala33 – open triangle; Arg34 – filled triangle). A schematic representation of the DNA template used for toeprinting is provided (RBS – ribosome binding site; NV1^5^ – sequence used to anneal the Yakima Yellow-labeled probe for reverse transcription).

**Supplementary Figure 3.**
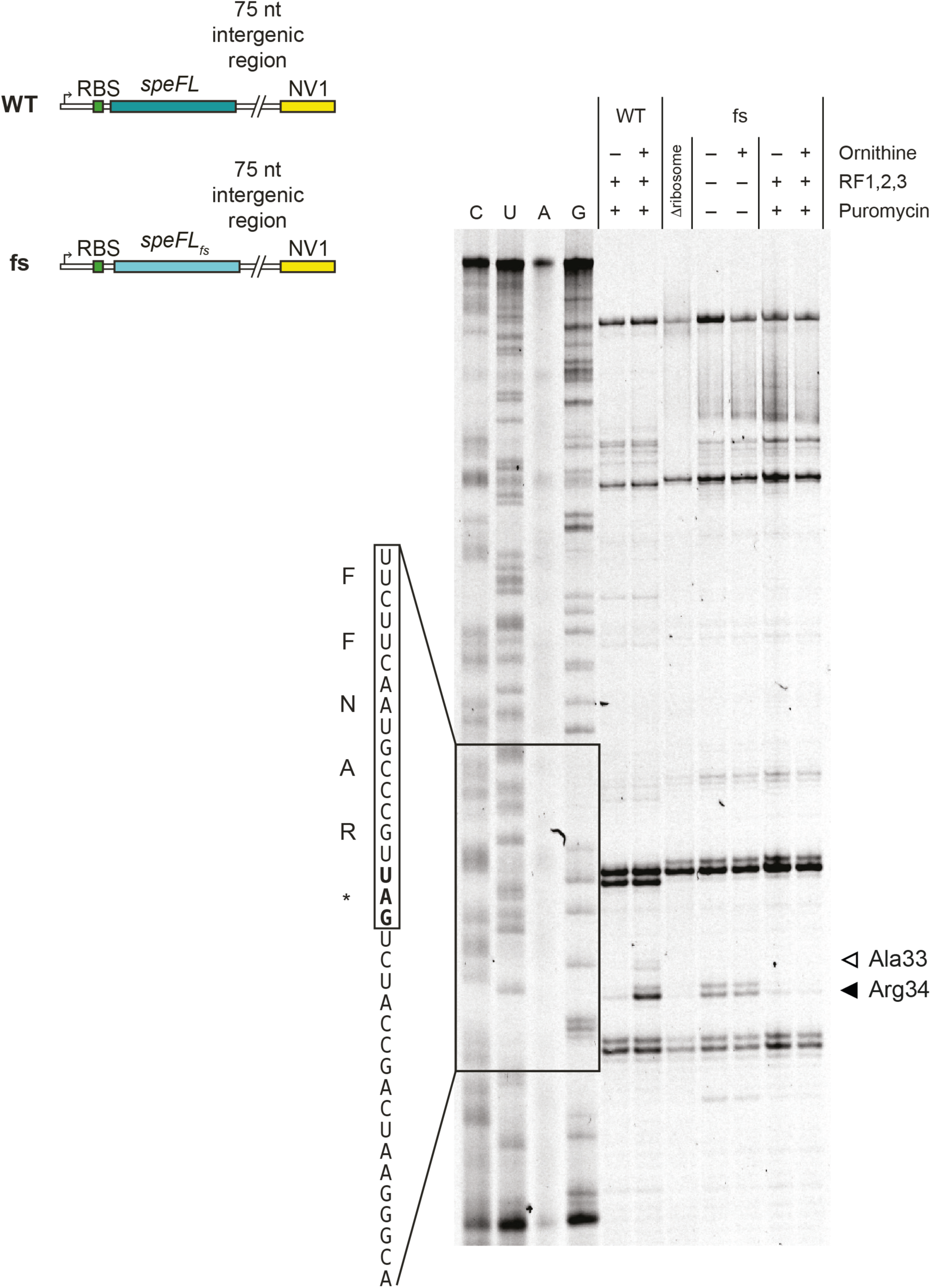
The amino acid sequence of SpeFL is important for ornithine-dependent translational arrest. Toeprinting assay^3, 4^ to monitor the translation of wild type (WT) and double frameshifted (*fs*) *speFL* in the presence (+) or absence (–) of 10 mM ornithine, release factors (RF1,2,3) or 90 μM puromycin. Arrows indicate ribosomes stalled with the codon for the indicated amino acid in the P-site (Ala33 – open triangle; Arg34 – filled triangle). Schematic representations of the DNA templates used for toeprinting are provided (RBS – ribosome binding site; NV1^5^ – sequence used to anneal the Yakima Yellow-labeled probe for reverse transcription).

**Supplementary Figure 4.**
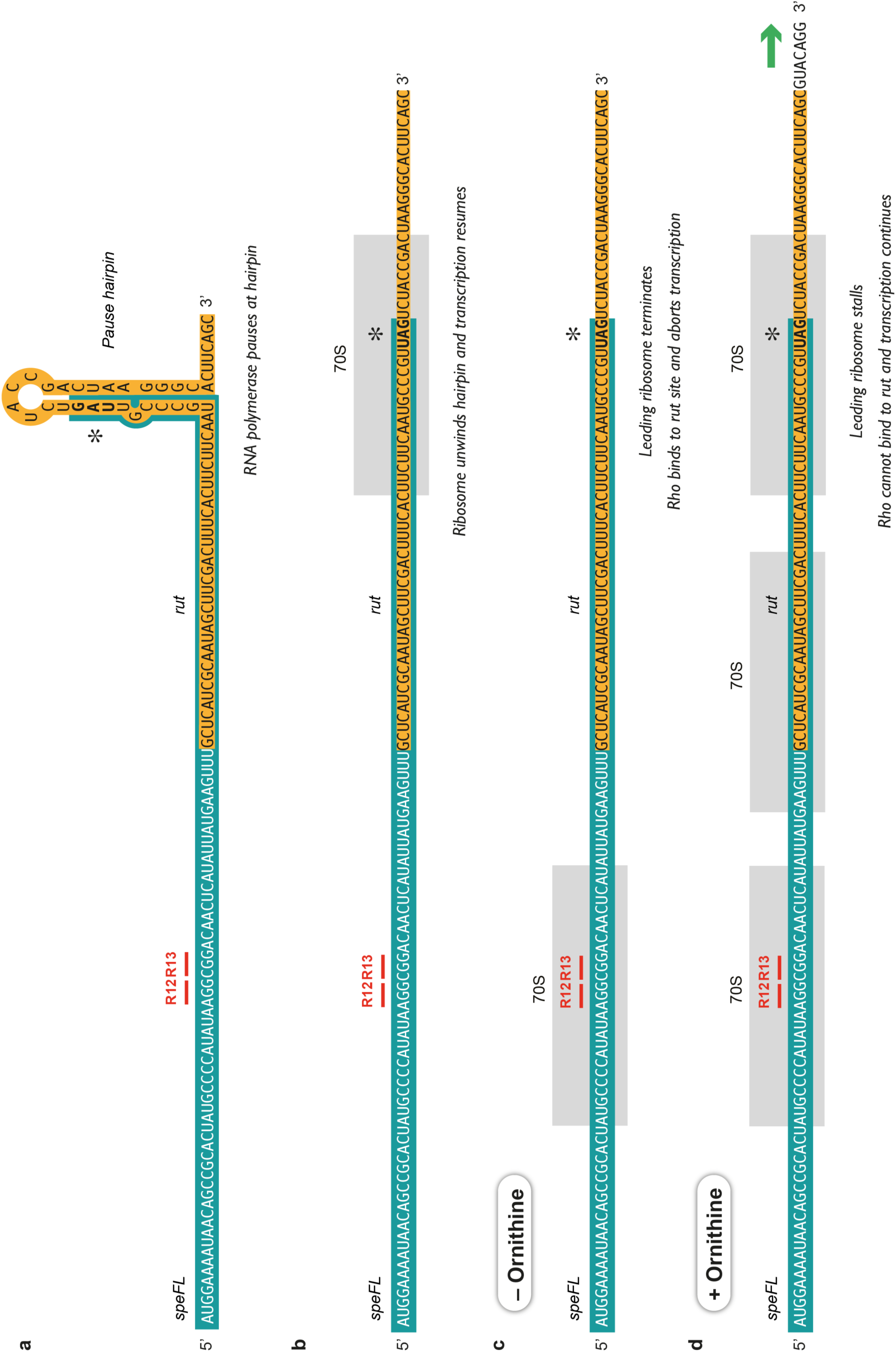
Mechanism of the SpeFL-and Rho-dependent regulation of the *speF* operon. The mRNA sequence of *speFL* and part of the adjacent intergenic region is shown at various stages of the induction process, namely (**a**) when the RNA polymerase pauses on a hairpin encompassing the 3’ end of *speFL*, (**b**) when the leading ribosome translating *speFL* unwinds the pause hairpin, (**c**) when the leading ribosome terminates translation in the absence of ornithine to allow Rho to bind to the *rut* site and (**d**) when the leading ribosome stalls in the presence of ornithine and blocks Rho binding, allowing the operon to be transcribed. The footprints of the ribosomes are in gray, *speFL* is in turquoise, the *rut* site is in yellow, rare codons R12 and R13 are in red and the UAG stop codon is indicated with an asterisk. The predicted 3’ end of the premature transcript is at position –1 of the consensus pause-inducing sequence element G_–11_G_–10_(C/T)_–1_G ^6^

**Supplementary Figure 5.**
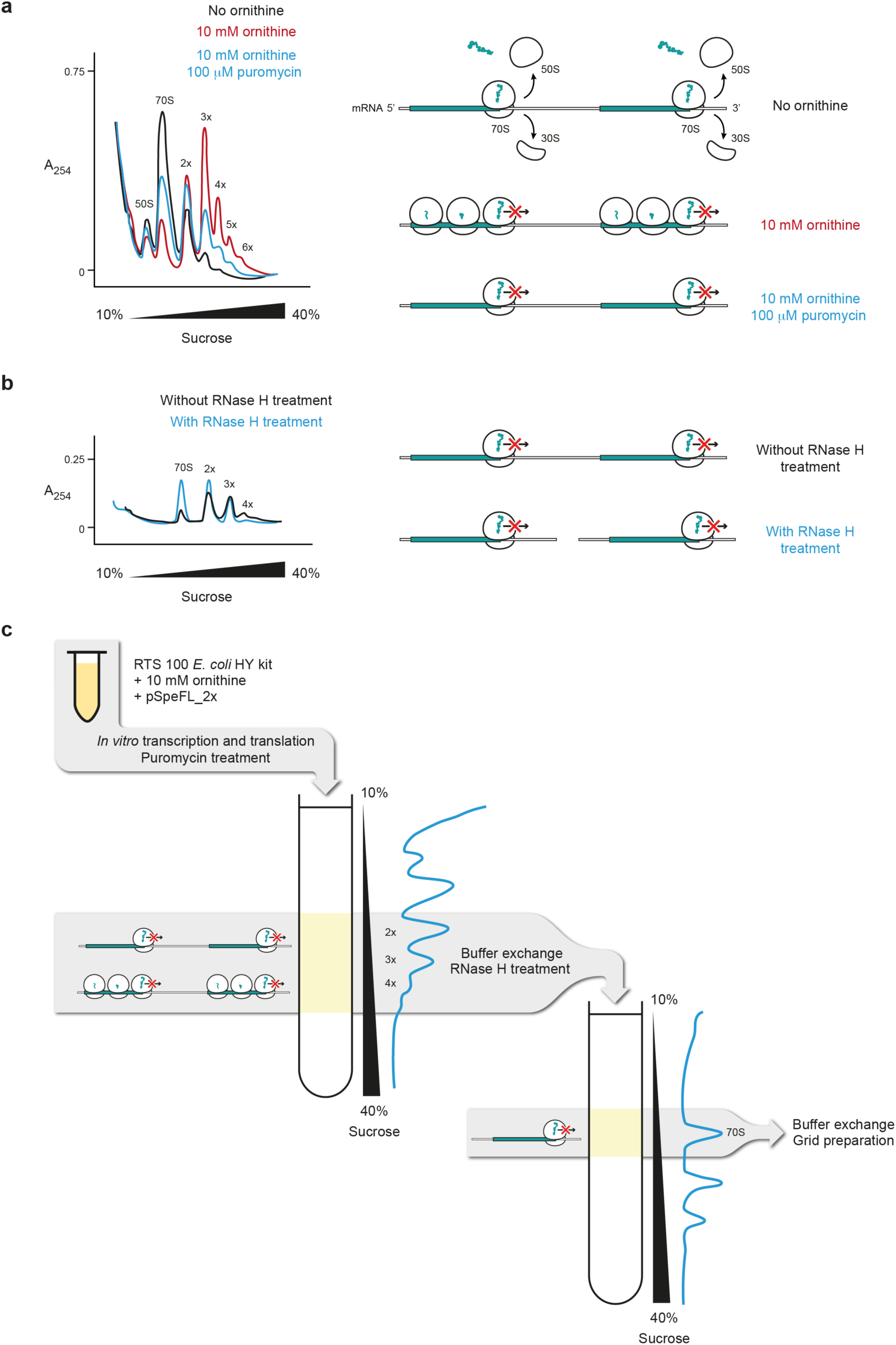
Purification of a SpeFL-70S complex stalled in the presence of ornithine. **a**, Overlaid absorbance profiles of sucrose gradients containing a translation mixture incubated without ornithine (black), in the presence of 10 mM L-ornithine (red) or in the presence of 10 mM L-ornithine followed by treatment with 100 µM puromycin (blue). A schematic diagram depicting the expected ribosomal species in each fraction is shown on the right. **b**, Overlaid absorbance profiles of sucrose gradients loaded with polysomal fractions from a, with (blue) or without (black) RNase H treatment. Expected ribosomal species for each fraction are shown on the right. **c**, Schematic representation of the purification strategy for SpeFL-70S. The collected fractions are indicated with gray boxes.

**Supplementary Figure 6.**
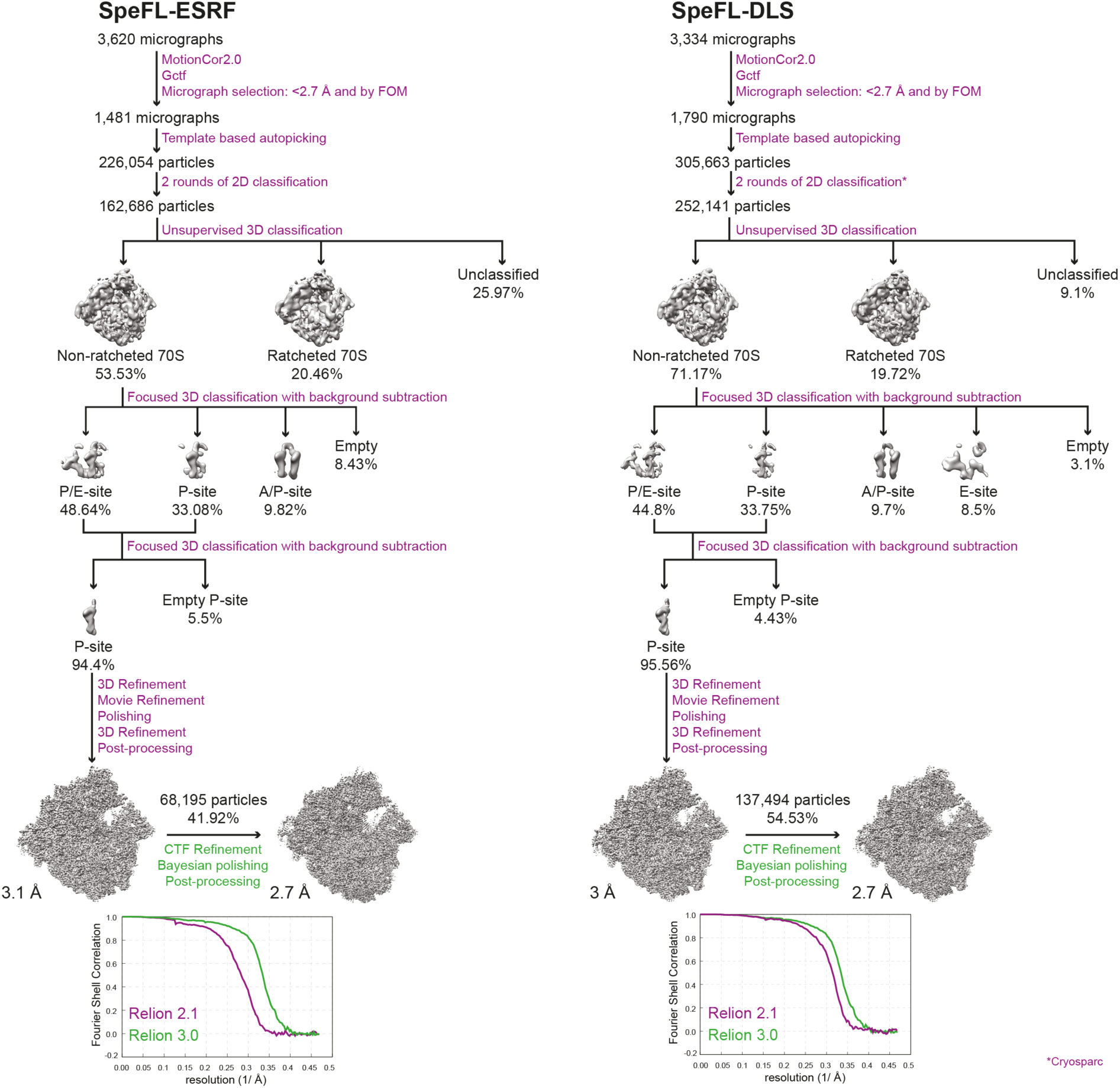
Flowchart of cryo-EM data processing for the SpeFL-ESRF and SpeFL-DLS datasets. Steps where *Relion 2.1*^7^ and *Relion 3.0*^8^ were used are shown in purple and green, respectively. The step where Cryosparc 0.6^9^ was used is indicated with an asterisk. Note the increase in resolution when using *Relion 3.0* compared to *Relion 2.1*. This increase was also matched by the quality of the resulting cryo-EM density. Both structures could be refined to an overall resolution of 2.7 Å using a Fourier shell correlation (FSC) cutoff of 0.143.

**Supplementary Figure 7.**
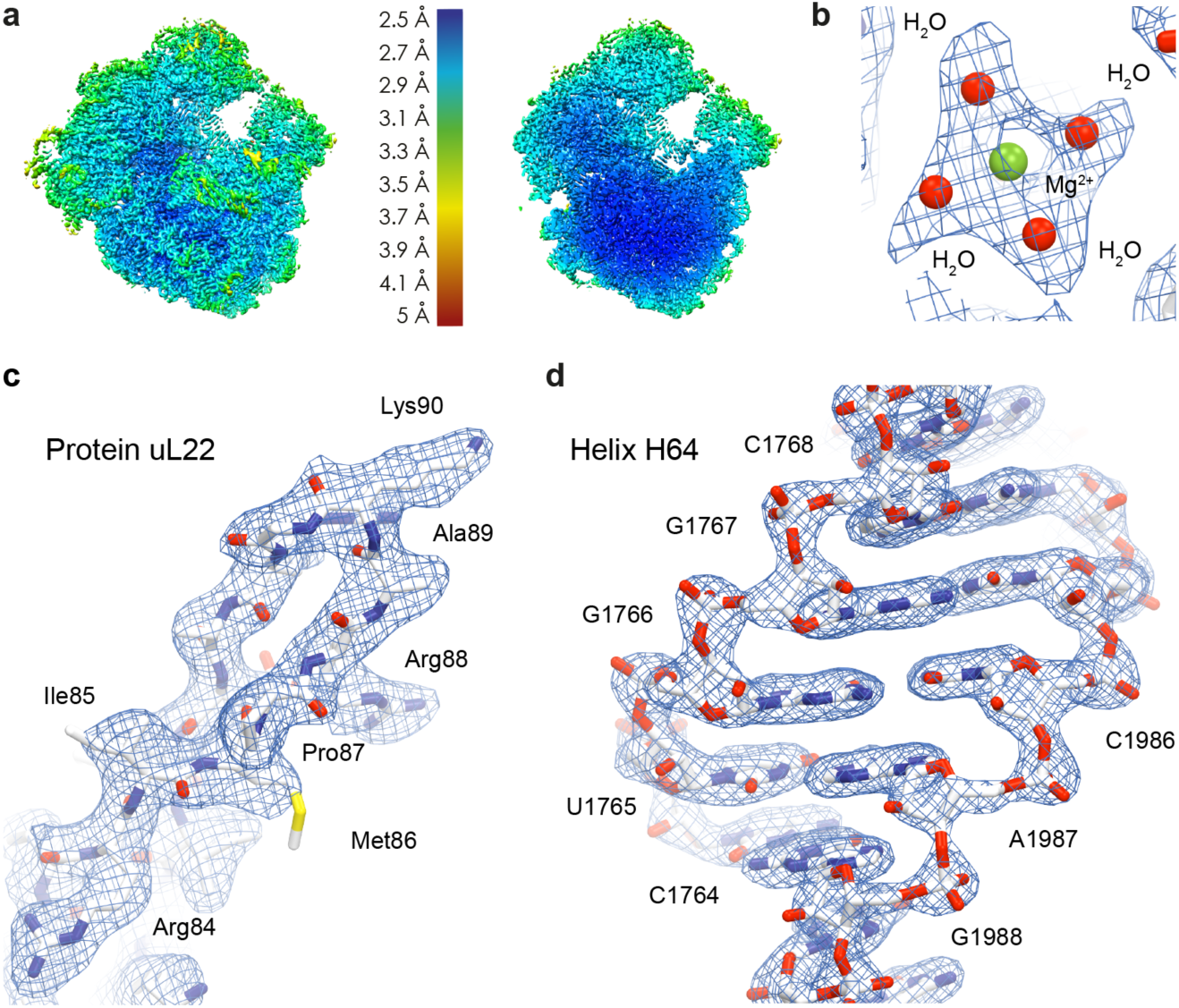
Quality of the cryo-EM reconstructions. **a**, Refined cryo-EM density map obtained in *Relion 3.0*^8^ filtered and colored by local resolution estimation values in *Chimera*^10^. A cross-section of the same map is also shown. **b,c,d,** Representative cryo-EM densities for (**b**) a hydrated magnesium ion bound to the 23S rRNA, (**c**) the tunnel extension of ribosomal protein uL22 and (**d**) helix H64 of the 23S rRNA.

**Supplementary Figure 8.**
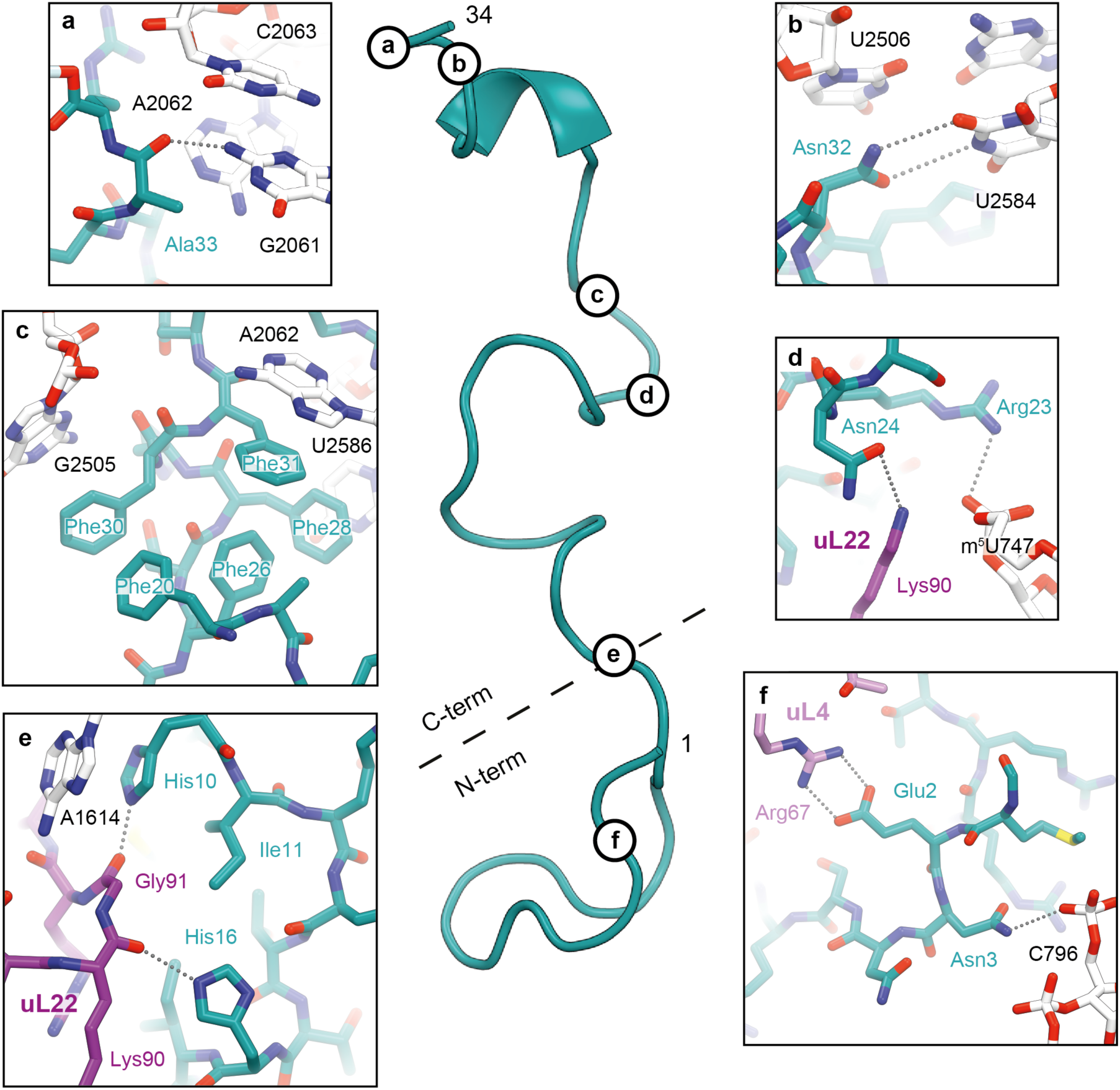
Interactions between SpeFL and the ribosome. A cartoon representation of SpeFL (turquoise) is shown in the middle panel. **a**, Potential hydrogen bond between Ala33 of SpeFL and the base of 23S rRNA residue G2061. **b**, Potential hydrogen bonds between Asn32 of SpeFL and the base of 23S rRNA residue U2584. **c**, Hydrophobic core of the SpeFL effector domain formed by residues Phe20, Phe26, Phe28, Phe30 and Phe31. Phe28, Phe30 and Phe31 of SpeFL form π-stacking interactions with the bases of 23S rRNA residues U2586, G2505 and A2062, respectively. **d**, Potential hydrogen bonds between Asn24 of SpeFL and Lys90 of ribosomal protein uL22, and electrostatic interaction between Arg23 of SpeFL and the phosphate backbone of 23S rRNA residue m^5^U747. **e**, The HIRRXXH ornithine-binding motif of SpeFL, showing potential hydrogen bonds between His10 and His16 of SpeFL, and Gly91 and Lys90 of ribosomal protein uL22, respectively. π-stacking interaction between 23S rRNA residue A1614 and His10 of SpeFL. **f**, Electrostatic interactions between residue Glu2 of SpeFL and residues Arg67 of ribosomal protein uL4. Possible hydrogen bond between residue Asn3 of SpeFL and the phosphate backbone of 23S rRNA residue C796.

**Supplementary Figure 9.**
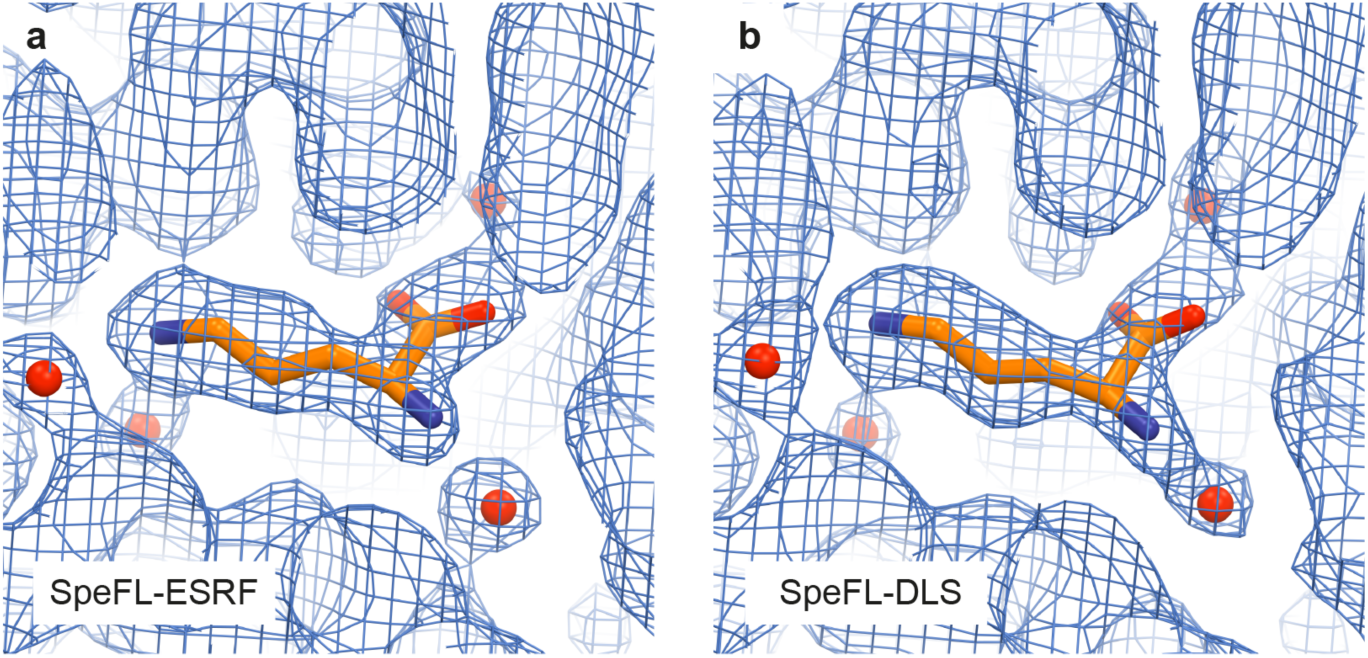
Sharpened cryo-EM density for L-ornithine and neighboring solvent molecules. A single L-ornithine molecule (orange) surrounded by 4 solvent molecules (red) is fitted into the cryo-EM density of the ligand binding pocket obtained for the (**a**) SpeFL-ESRF and (**b**) SpeFL-DLS datasets. Note that peaks for the solvent molecules are visible in the two independently determined cryo-EM maps, indicating that these densities cannot be attributed to random noise.

**Supplementary Figure 10.**
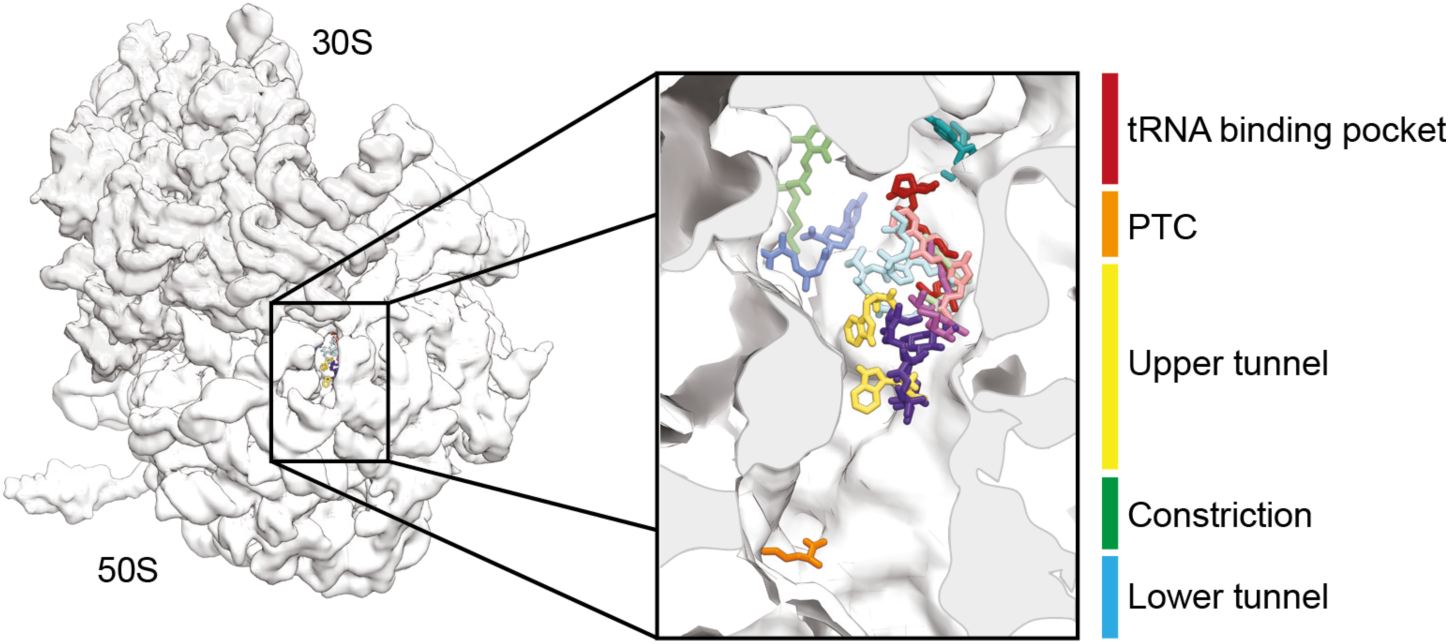
Small ligand binding pockets of the ribosomal exit tunnel. Overview and close-up view of a cross-section of the *E. coli* 70S ribosomal exit tunnel showing the L-ornithine molecule observed in this work (orange) together with small molecules that are known to bind to the ribosomal exit tunnel: blasticin S (PDB: 4v9q, dark blue)^11^, chloramphenicol (PDB: 4v7w, light green)^12^, clindamycin (PDB: 4v7v, magenta)^13^, dalfopristin (PDB: 4u24, light blue)^14^, erythromycin (PDB: 4v7u, purple)^13^, hygromycin (PDB: 5dox, red)^15^, linezolid (PDB: 3dll, pink)^16^, puromycin (PDB: 1q82, cyan)^17^, sparsomycin (PDB: 1njn, dark green)^18^ and tryptophan (PDB: 4uy8, yellow)^19^. The different regions of the tunnel are highlighted: the tRNA binding pocket (red), the peptidyl transferase center (PTC) (orange), the upper tunnel (yellow), the constriction formed by uL22 and uL4 (green) and the lower tunnel (blue).

**Supplementary Figure 11.**
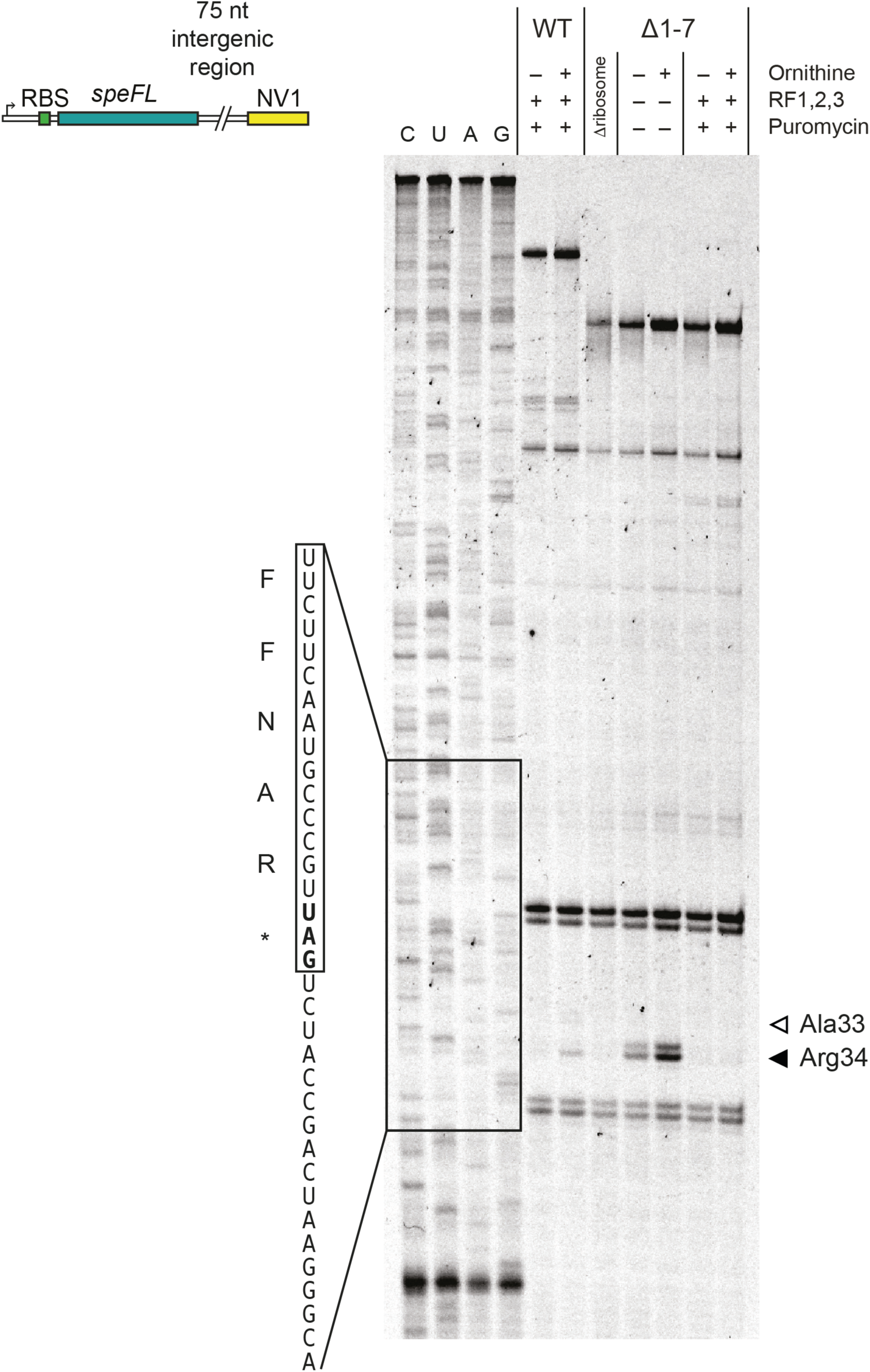
Importance of residues 1–7of SpeFL. Toeprinting assay^3, 4^ to monitor the translation of wild-type *speFL* and *speFL*Δ1–7 in the absence (–) or presence (+) of 10 mM ornithine, release factors (RF1,2,3) or 90 μM puromycin. Arrows indicate ribosomes stalled with the indicated amino acid in the P-site (Ala33 – open triangle; Arg34 – filled triangle). A schematic representation of the DNA template used for toeprinting is provided (RBS – ribosome binding site; NV1^5^ – sequence used to anneal the Yakima Yellow-labeled probe for reverse transcription).

**Supplementary Figure 12.**
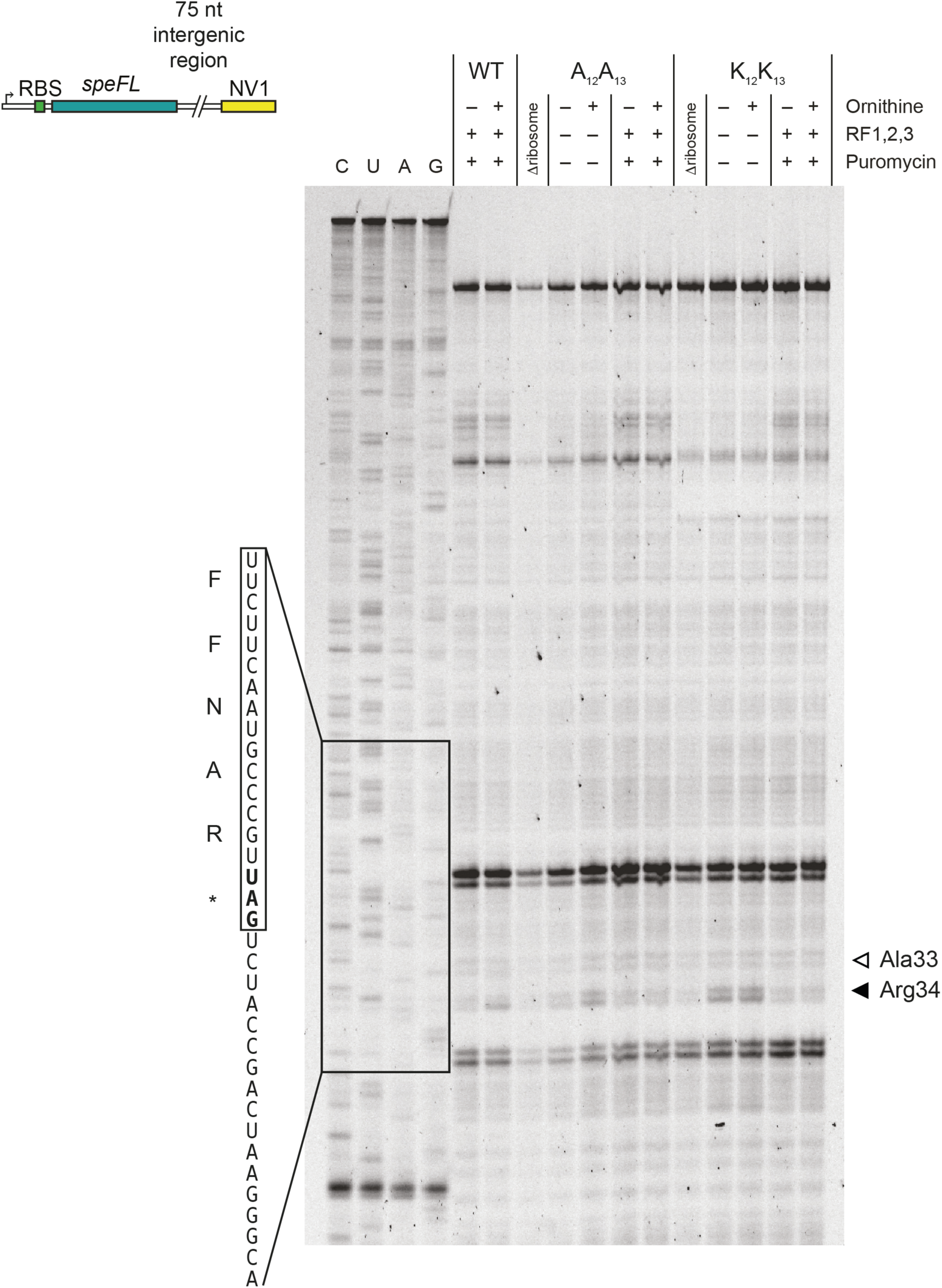
Importance of residues 12 and 13 of SpeFL. Toeprinting assay^3, 4^ to monitor the translation of wild-type *speFL, speFL-R12A-R13A* (A_12_A_13_) and *speFL-R12K-R13K* (K_12_K_13_) in the absence (–) or presence (+) of 10 mM ornithine, release factors (RF1,2,3) or 90 μM puromycin. Arrows indicate ribosomes stalled with the indicated amino acid in the P-site (Ala33 – open triangle; Arg34 – filled triangle). A schematic representation of the DNA template used for toeprinting is provided (RBS – ribosome binding site; NV1^5^ – sequence used to anneal the Yakima Yellow-labeled probe for reverse transcription).

**Supplementary Figure 13.**
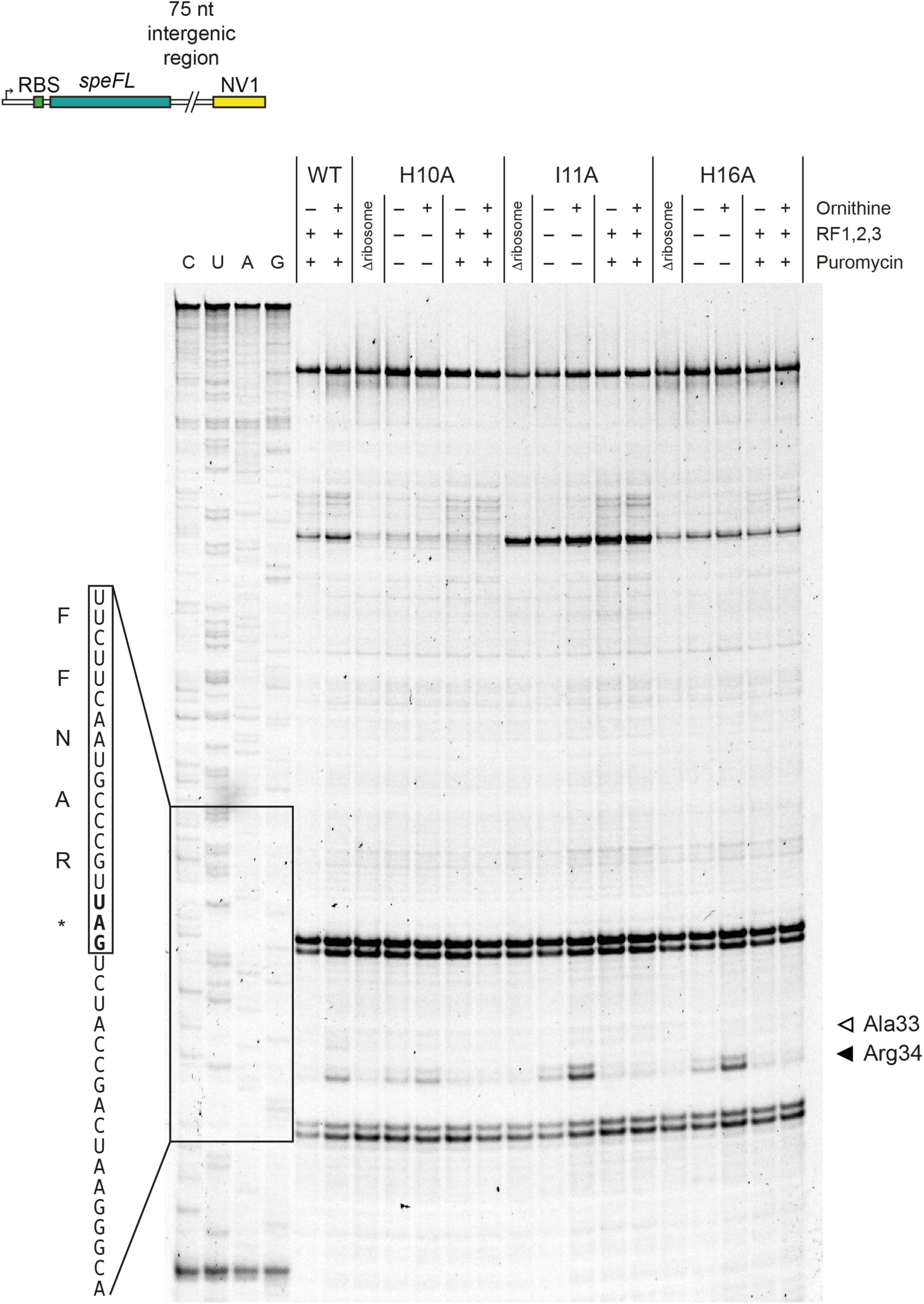
Importance of residues 10, 11 and 16 of SpeFL. Toeprinting assay^3, 4^ to monitor the translation of wild-type *speFL, speFL-H10A, speFL-I11A* and *speFL-H16A* in the absence (–) or presence (+) of 10 mM ornithine, release factors (RF1,2,3) or 90 μM puromycin. Arrows indicate ribosomes stalled with the indicated amino acid in the P-site (Ala33 – open triangle; Arg34 – filled triangle). A schematic representation of the DNA template used for toeprinting is provided (RBS – ribosome binding site; NV1^5^ – sequence used to anneal the Yakima Yellow-labeled probe for reverse transcription).

**Supplementary Table 1.**
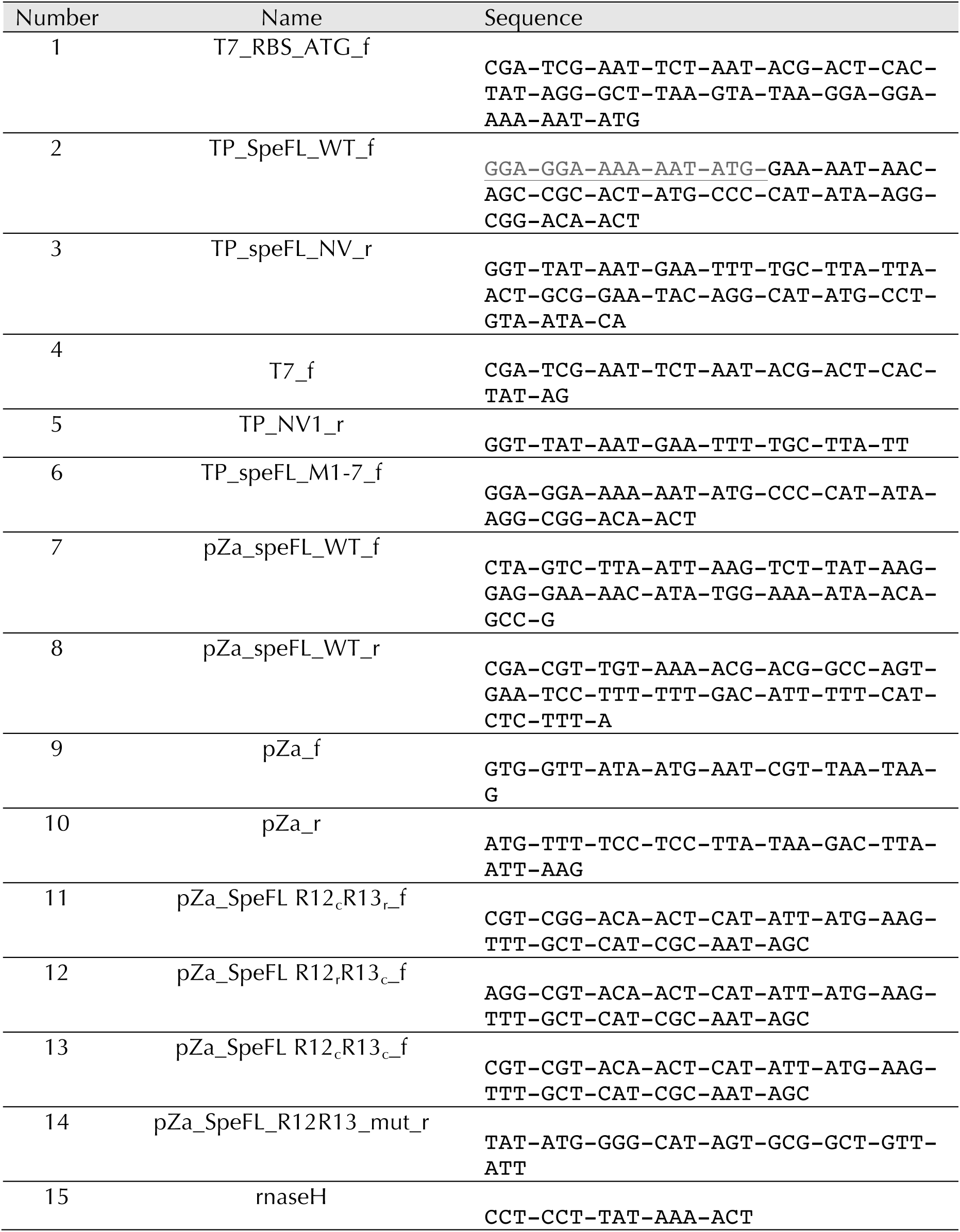
Oligonucleotides

**Supplementary Table 2.**
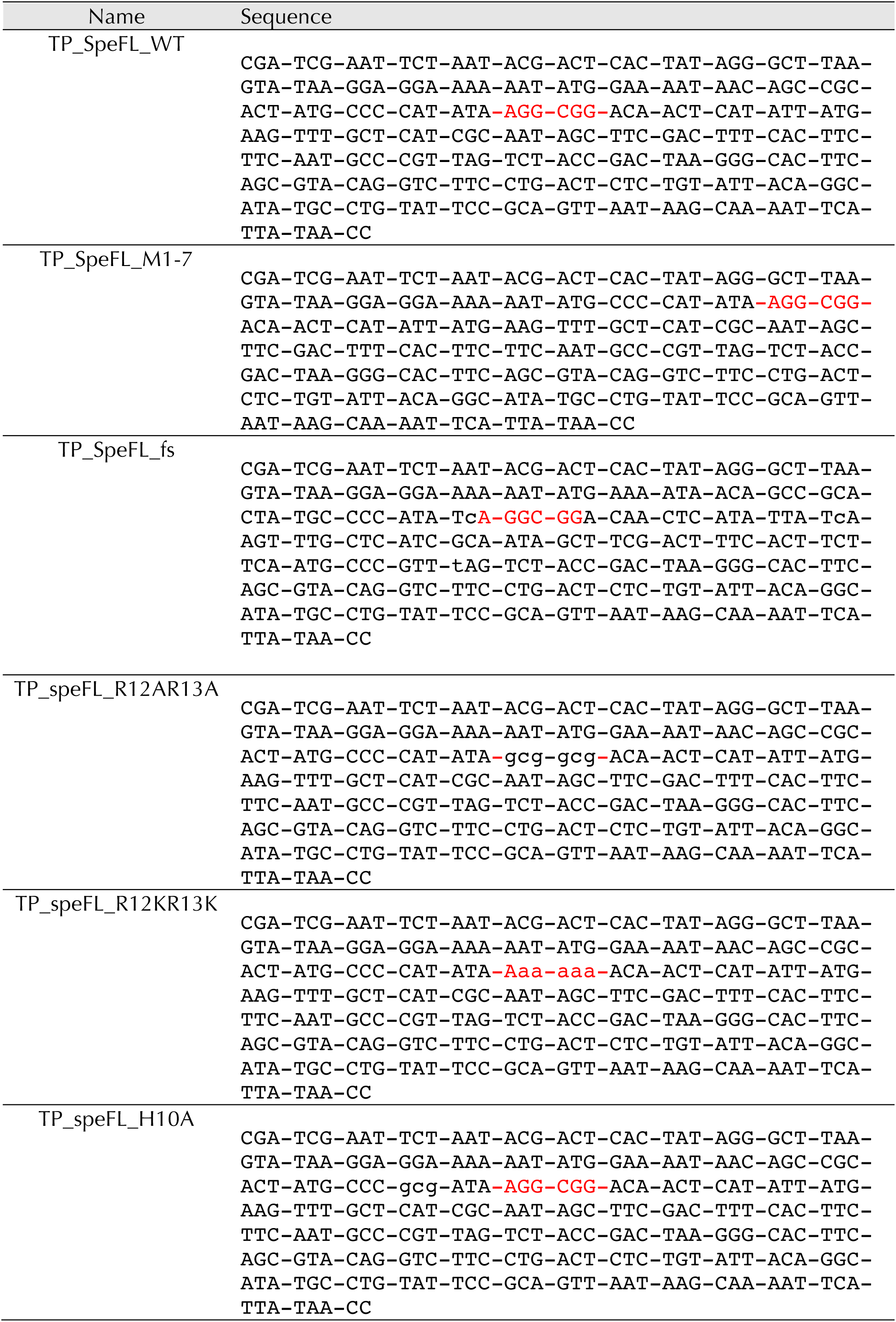
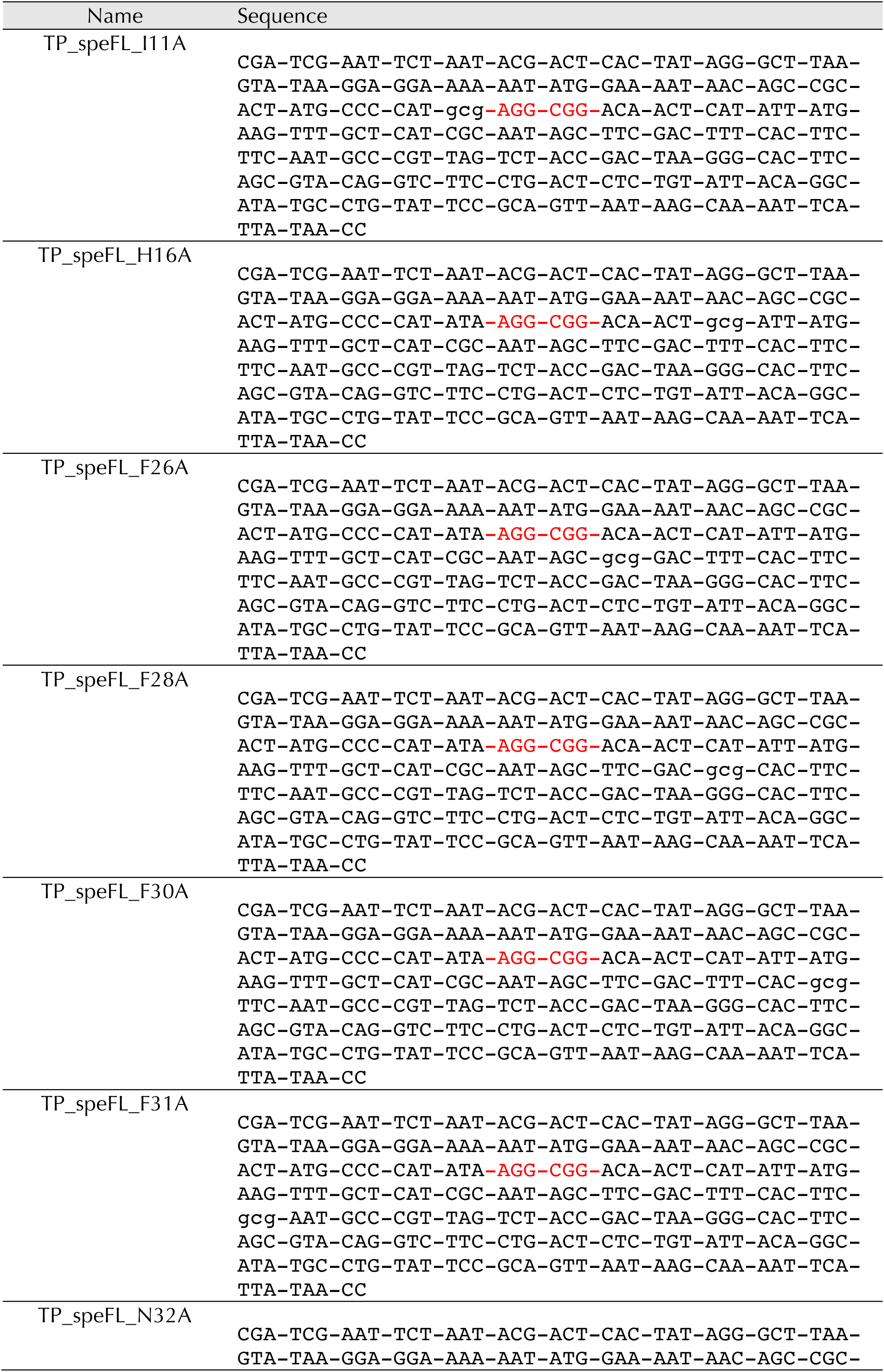
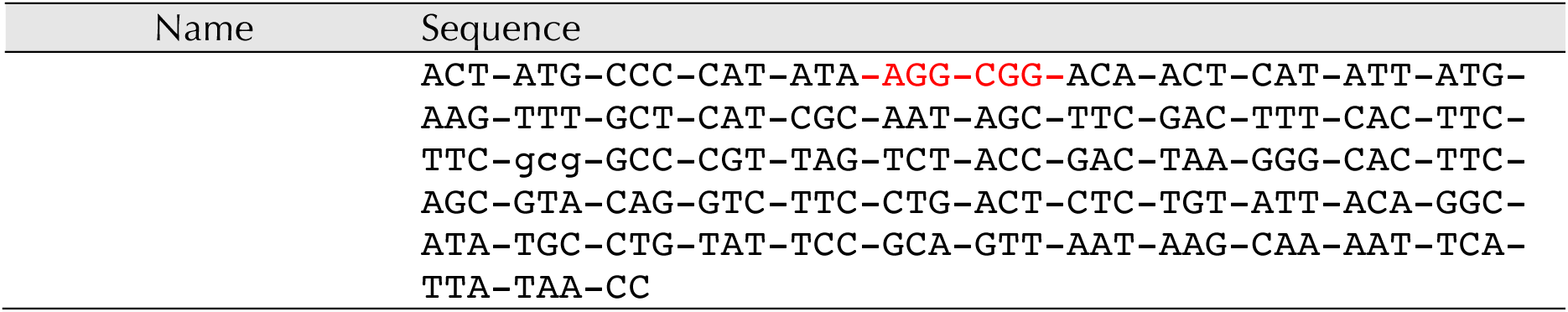
DNA templates

**Supplementary Table 3.** Cryo-EM statistics and model refinement

|  | #1 SpeFL-ESRF<br>(EMDB-10458)<br>(PDB 6TC3) | #2 SpeFL-DLS<br>(EMDB-10453)<br>(PDB 6TBV) |
| --- | --- | --- |
| **Data collection and processing** |  |  |
| Magnification | 130,000x | 130,000x |
| Voltage (kV) | 300 | 300 |
| Electron exposure (e-/Å<sup>2</sup>) | 30 | 29.6 |
| Defocus range (μm) | -0.5 to -1.6 | -0.6 to -1.5 |
| Pixel size (Å) | 1.067 | 1.067 |
| Symmetry imposed | C1 | C1 |
| Initial particle images (no.) | 226,054 | 305,663 |
| Final particle images (no.) | 68,195 | 137,494 |
| Map resolution (Å) | 2.7 | 2.7 |
| FSC threshold | 0.143 | 0.143 |
| Map resolution range (Å) | 2.5-10 | 2.5-8.7 |
| **Refinement** |  |  |
| Initial model used (PDB code) | 4ybb | 4ybb |
| Model resolution (Å) | 2.1 | 2.1 |
| FSC threshold | 0.143 | 0.143 |
| Model resolution range (Å) | 2.1-69.39 | 2.1-69.39 |
| Map sharpening *B* factor (Å<sup>2</sup>) | -10 | -10 |
| Model composition |  |  |
| Non-hydrogen atoms | 245,116 | 245,116 |
| Protein residues | 10,310 | 10,310 |
| Ligands |  |  |
| *B* factors (Å<sup>2</sup>) |  |  |
| Protein | 48.22 | 53.05 |
| Ligand | 34.32 | 38.14 |
| R.m.s. deviations |  |  |
| Bond lengths (Å) | 0.01 | 0.01 |
| Bond angles (°) | 0.97 | 0.93 |
| Validation |  |  |
| MolProbity score | 1.21 | 1.17 |
| Clashscore | 1.82 | 1.66 |
| Poor rotamers (%) | 0.41 | 0.6 |
| Ramachandran plot |  |  |
| Favored (%) | 96.12 | 96.27 |
| Allowed (%) | 3.74 | 3.66 |
| Disallowed (%) | 0.14 | 0.07 |

**Supplementary Data 1.**
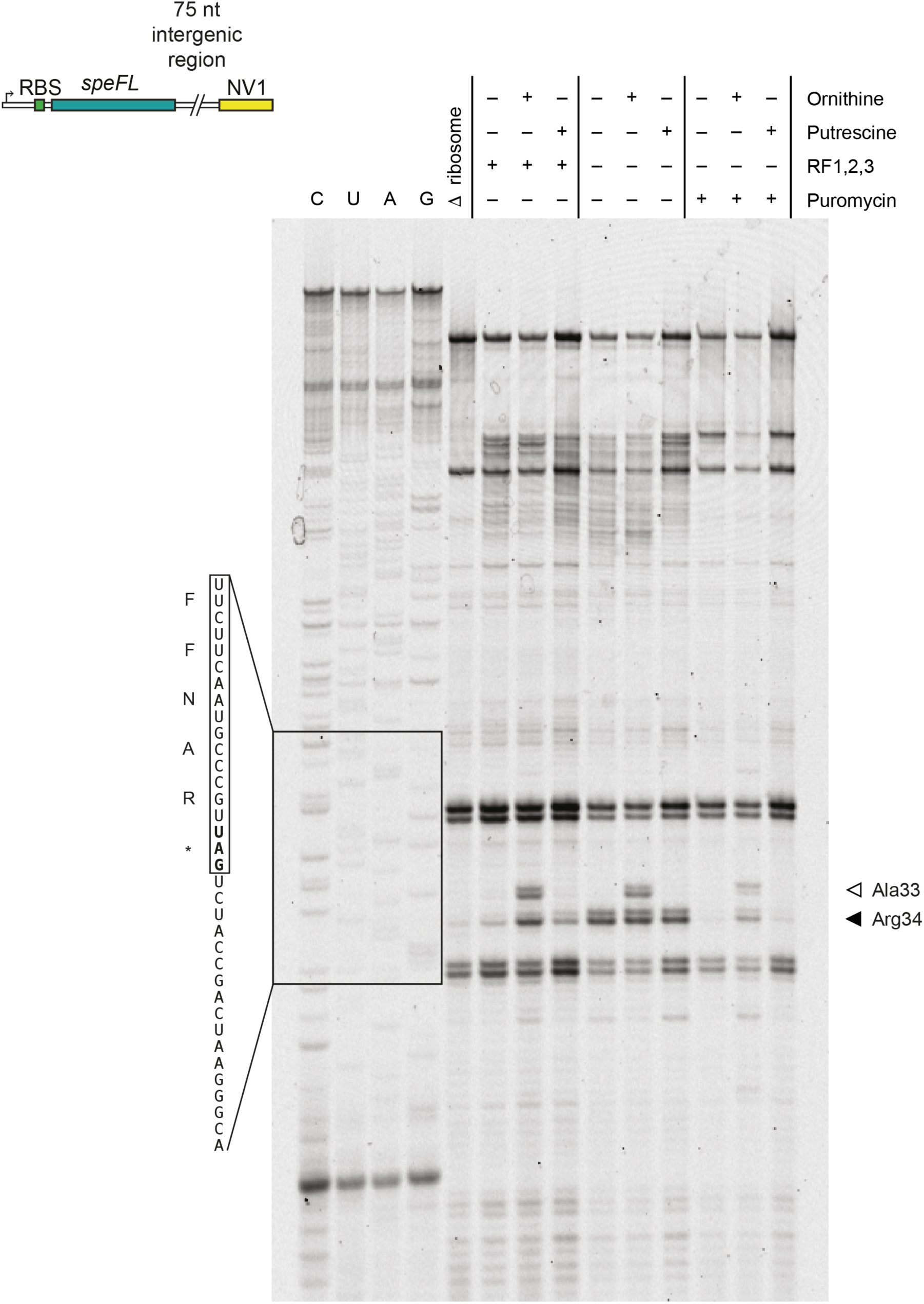
Ornithine-dependent ribosomal stalling on speFL. Toeprinting assay^3, 4^ to monitor the translation of *speFL* in the absence (–) or presence (+) of 10 mM ornithine, 10 mM putrescine, release factors (RF1,2,3) or 90 μM puromycin. Arrows indicate ribosomes stalled with the codon for the indicated amino acid in the P-site (Ala33 – open triangle; Arg34 – filled triangle). A schematic representation of the DNA template used for toeprinting is provided (RBS – ribosome binding site; NV1^5^ – sequence used to anneal the Yakima Yellow-labeled probe for reverse transcription).

**Supplementary Data 2.**
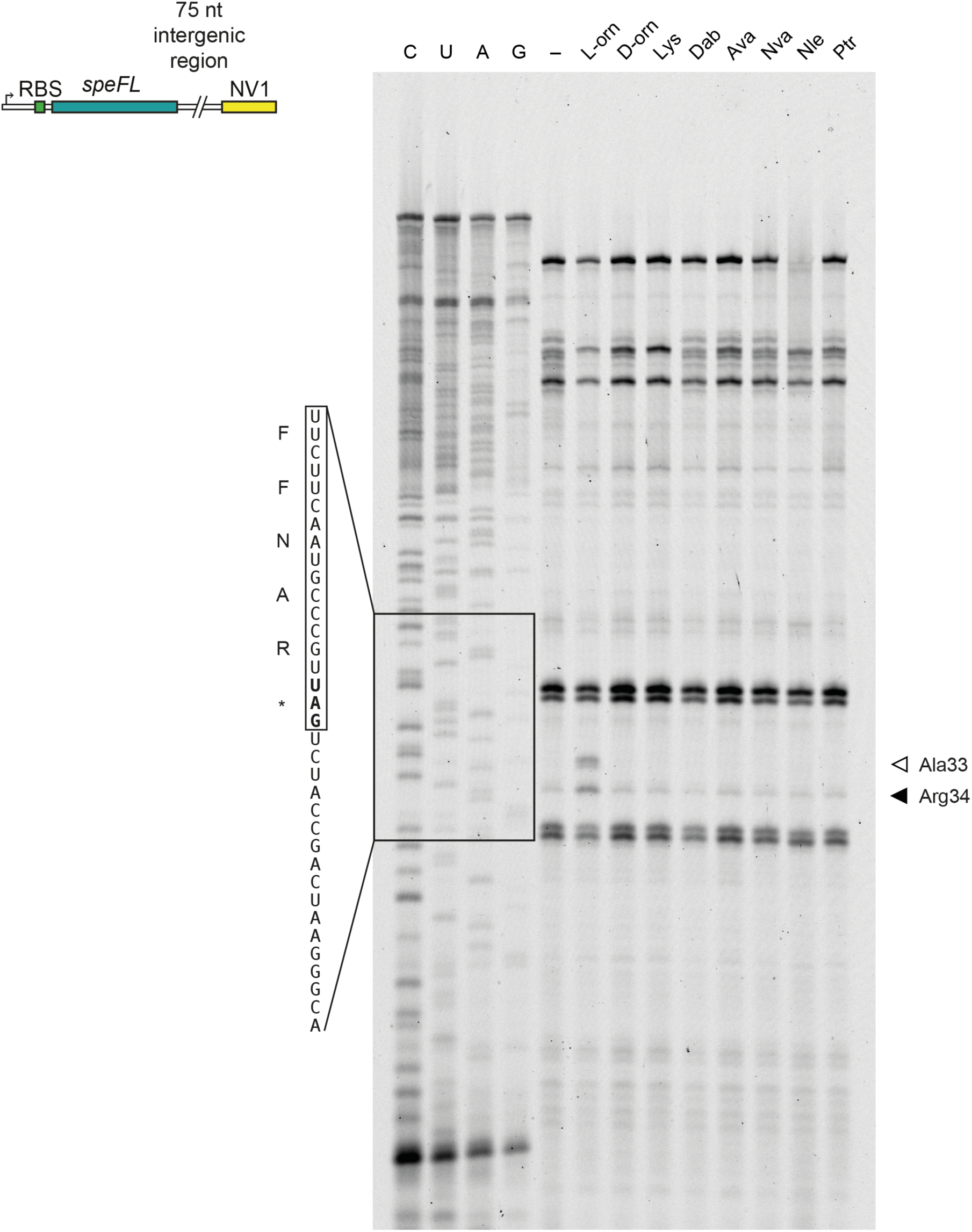
Selectivity of SpeFL for L-ornithine. Toeprinting assay^3, 4^ to monitor the translation of wild-type (WT) *speFL* in the absence (–) or presence of 10 mM (+) of various small molecules (see Fig. 2e for details). All samples were treated with 90 µM puromycin. Arrows indicate ribosomes stalled with the indicated amino acid in the P-site (Ala33 – open triangle; Arg34 – filled triangle). A schematic representation of the DNA template used for toeprinting is provided (RBS – ribosome binding site; NV1^5^ – sequence used to anneal the Yakima Yellow-labeled probe for reverse transcription).

**Supplementary Data 3.**
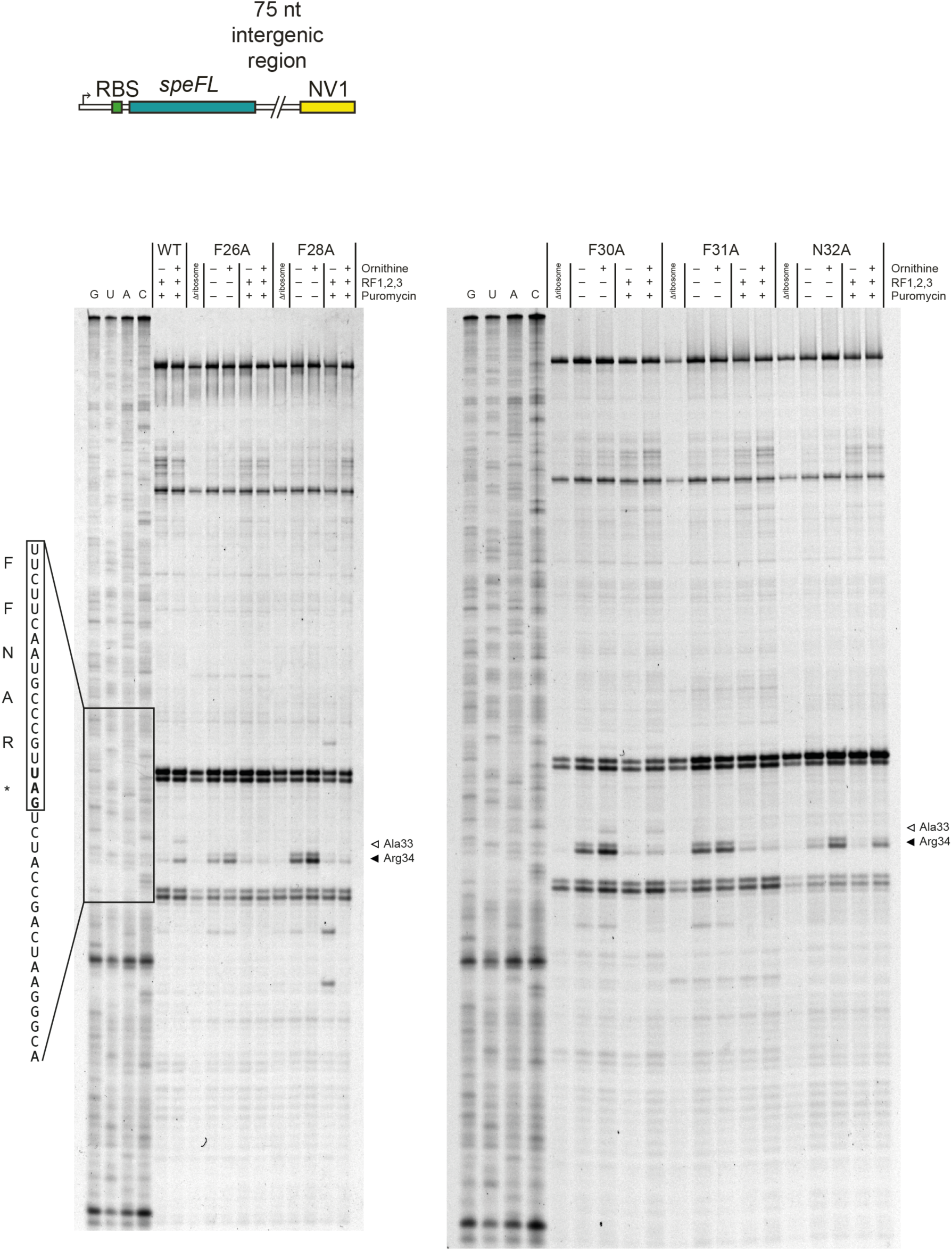
Aromatic residues in the SpeFL effector domain are important for translational arrest. Toeprinting assay^3, 4^ to monitor the translation of wild-type (WT) and mutant *speFL* in the absence (–) or presence (+) of 10 mM ornithine, release factors (RF1,2,3) or 90 μM puromycin. Arrows indicate ribosomes stalled with the indicated amino acid in the P-site (Ala33 – open triangle; Arg34 – filled triangle). A schematic representation of the DNA template used for toeprinting is provided (RBS – ribosome binding site; NV1^5^ – sequence used to anneal the Yakima Yellow-labeled probe for reverse transcription).

